# Direct interrogation of context-dependent GPCR activity with a universal biosensor platform

**DOI:** 10.1101/2024.01.02.573921

**Authors:** Remi Janicot, Marcin Maziarz, Jong-Chan Park, Alex Luebbers, Elena Green, Jingyi Zhao, Clementine Philibert, Hao Zhang, Mathew D. Layne, Joseph C. Wu, Mikel Garcia-Marcos

**Affiliations:** Department of Biochemistry & Cell Biology, Chobanian & Avedisian School of Medicine, Boston University, Boston, MA 02118, USA; Stanford Cardiovascular Institute, Stanford University School of Medicine, Stanford, CA 94305, USA; Department of Biology, College of Arts & Sciences, Boston University, Boston, MA 02115, USA

## Abstract

G protein-coupled receptors (GPCRs) are the largest family of druggable proteins in the human genome, but progress in understanding and targeting them is hindered by the lack of tools to reliably measure their nuanced behavior in physiologically-relevant contexts. Here, we developed a collection of compact <u>ONE</u> vector <u>G</u>-protein <u>O</u>ptical (ONE-GO) biosensor constructs as a scalable platform that can be conveniently deployed to measure G-protein activation by virtually any GPCR with high fidelity even when expressed endogenously in primary cells. By characterizing dozens of GPCRs across many cell types like primary cardiovascular cells or neurons, we revealed new insights into the molecular basis for G-protein coupling selectivity of GPCRs, pharmacogenomic profiles of anti-psychotics on naturally-occurring GPCR variants, and G-protein subtype signaling bias by endogenous GPCRs depending on cell type or upon inducing disease-like states. In summary, this open-source platform makes the direct interrogation of context-dependent GPCR activity broadly accessible.

## INTRODUCTION

G-protein-coupled receptors (GPCRs) mediate responses to many natural stimuli, including most neurotransmitters and two-thirds of hormones^1–3^, and are also the target for 30-40% of clinically-approved drugs^4,5^. While representing the largest family of druggable targets in the human genome, there is still vast untapped potential because the majority of GPCRs remain understudied^6^. With the advent of wide-spread success in elucidating high resolution structures of many GPCRs^7,8^, current trends in GPCR research and drug development are focused on identifying mechanisms and compounds that fine tune signaling outcomes, including allosteric modulation or ligand-directed biased signaling^9–12^. However, progress in this area has been historically hampered by the overreliance on assays that measure GPCR activity indirectly (e.g., second messengers), and on a limited number of cells lines, which do not necessarily recapitulate GPCR physiological contexts. As a consequence, pharmacological features of new compounds characterized by *in vitro* functional assays of GPCR activity may not accurately translate to the expected therapeutic benefit *in vivo*^13^, as illustrated by ongoing controversies on novel opioid drugs with diminished side effects^14–19^. A critical factor for the discrepancy between *in vitro* and *in vivo* pharmacology is the existence of *system biased GPCR signaling*, i.e., how signaling pathways are hardwired in each specific cellular system to propagate the initial drug-receptor interaction to a cellular response^13,20^.

An approach to mitigate the impact of system bias in drug discovery is the use of assays that measure GPCR activity more directly to avoid potential crosstalk and amplification mechanisms specific to a given cellular system. In this regard, GPCRs propagate signaling primarily by directly activating heterotrimeric G proteins (Gαβγ), although some responses are mediated by β-arrestins^1,2,21^. GPCRs activate G proteins by promoting the exchange of GDP for GTP on Gα subunits. In turn, this leads to the dissociation of Gα-GTP and free Gβγ^22,23^. The latter event has been widely exploited to generate biosensors of G protein activity based on fluorescence or bioluminescence resonance energy transfer (FRET or BRET, respectively)^24–28^. While Gα/Gβγ dissociation is an event proximal to GPCR activation, these approaches are not broadly applicable to physiologically relevant systems and can significantly compromise the fidelity of the responses measured. For example, even in optimized versions of biosensors that measure Gα/Gβγ dissociation (e.g., TRUPATH^29^), overexpression of exogenous G protein subunits tagged with BRET donors and acceptors is required. This imposes several limitations. One is that multiple genetic components must be delivered simultaneously to cells in optimal proportions, making the approach only feasible in a few easily transfectable cell lines. Another major limitation is that exogenous G protein subunits can form non-BRET productive complexes with endogenous G proteins, so they must be expressed in large excess to yield robust BRET responses. This represents a significant perturbation of the system under investigation by the method of detection employed.

We recently developed alternative G protein biosensor designs to measure levels of Gα-GTP in cells as a more proximal event to receptor activation than Gα/Gβγ dissociation and converted them into single polypeptide chain biosensors named BERKY^30^. While this allowed the detection of activation of endogenous GPCRs in primary neurons and without interfering with downstream signaling^30^, the small dynamic range has remained a limitation for the broad applicability of this type of biosensor. Others have developed an effector membrane translocation assay (EMTA) that also measures Gα-GTP^31^. However, some EMTA biosensors are likely to interfere with G protein signaling, and the broad applicability of EMTA with endogenous GPCRs in primary cells has not been established, probably because of the need to deliver simultaneously multiple genetic components. A notable exception for both BERKY and EMTA biosensor systems is the lack of direct detection of Gα-GTP species for one of the four G protein families, G_s_^22^.

In the work presented here, we set out to generate a comprehensive collection of Gα-GTP biosensor constructs for different G protein types, including the evasive Gαs-GTP, that combined large dynamic range of detection with the ability to monitor endogenous GPCR activity across a wide range of physiologically-relevant systems (e.g., different primary cell types). By reducing the number of biosensor components and condensing them into a single vector that expresses the BRET pairs in appropriate proportions, we achieved a system that can be deployed with ease to scale up throughput and to permit the investigation of endogenous GPCRs in diverse cell types. To facilitate adoption and customization by others, all the biosensor constructs described here and their designs have been made publicly accessible through Addgene. Because the system relies on assembling functional G protein complexes between endogenous Gβγ and low levels of exogenous Gα, the interference with endogenous signaling is imperceptible. By leveraging the advantages of this biosensor platform, we reveal that endogenous GPCRs in primary cells display unique signaling behaviors depending on the cellular context, raising the important point that system bias must be considered even when measuring the most direct readout of GPCR signal transduction (i.e., nucleotide exchange on Gα).

## RESULTS

### Direct detection of active Gαs in cells with a BRET biosensor

We sought to identify a protein sequence that would bind specifically to active Gαs, anticipating that it would serve as a specific detector module for a genetically encoded live-cell biosensor. Initial efforts based on previously described Gαs interactors^32–34^ failed to identify a suitable detector module (*not shown*). We turned our attention to the patent literature and found a series of putative Gαs-GTP binding peptides from a phage display screen (**Fig. 1A**^35^). We tested 12 of these peptides for binding to active Gαs (**Fig. 1A**) and determined that sequences KB1691 and KB2123 bound strongly to active Gαs, but not active Gαo (**Fig. 1A**). To test whether KB1691 or KB2123 could serve as detector modules in BRET-based biosensors, they were fused to membrane anchored nanoluciferase (Nluc, BRET donor). This plasmids was co-expressed in HEK293T with Venus-tagged Gαs (BRET acceptor), along with Gβ_1_, Gγ_2_, and the β2 adrenergic receptor (β2AR) (**Fig. 1B, Fig. S1**). We systematically tested three versions of Gαs (short isoform) in which Venus was inserted at different internal locations^36^, with and without co-expression of the Gαs chaperone Ric-8B^37^. Both KB1691- and KB2123-based sensors performed best with the Gαs constructs with Venus inserted at codon 99 (Gαs-99V) and in the presence of Ric-8B— i.e., cells expressing these components led to larger BRET increases upon stimulation with the β-adrenergic agonist isoproterenol, which were reversed upon treatment with the antagonist alprenolol (**Fig. 1B, Fig. S1**). Similar results were obtained with a second GPCR known to activate Gαs, the dopamine D1 receptor (D1R) (**Fig. S1C**). Amino acid position 99 is in a loop of the helical domain of Gαs that tolerates insertion of bulky tags, as previously demonstrated by the functional validation of Gαs-99V and other constructs with insertions in the same loop^25,38^. While Ric-8B expression improved the magnitude of the responses detected, this was not due to a significant increase in Gαs-99V expression (**Fig. S1**). This suggests that the improved responses are due to better folding and/or membrane targeting of tagged Gαs, as previously reported for Ric-8 proteins by others^37^. An alternative biosensor design relying on bystander BRET^39,40^ with untagged Gαs also yielded responses (**Fig. S2**), but was not as robust as the membrane-anchored, Nluc-fused peptides and Gαs-99V in the presence of Ric-8B, which became the biosensor design of choice for subsequent studies. These results indicate that GPCR-mediated activation of Gαs can be detected in cells with an optical biosensor based on peptides that bind Gαs-GTP.

**Figure 1.**
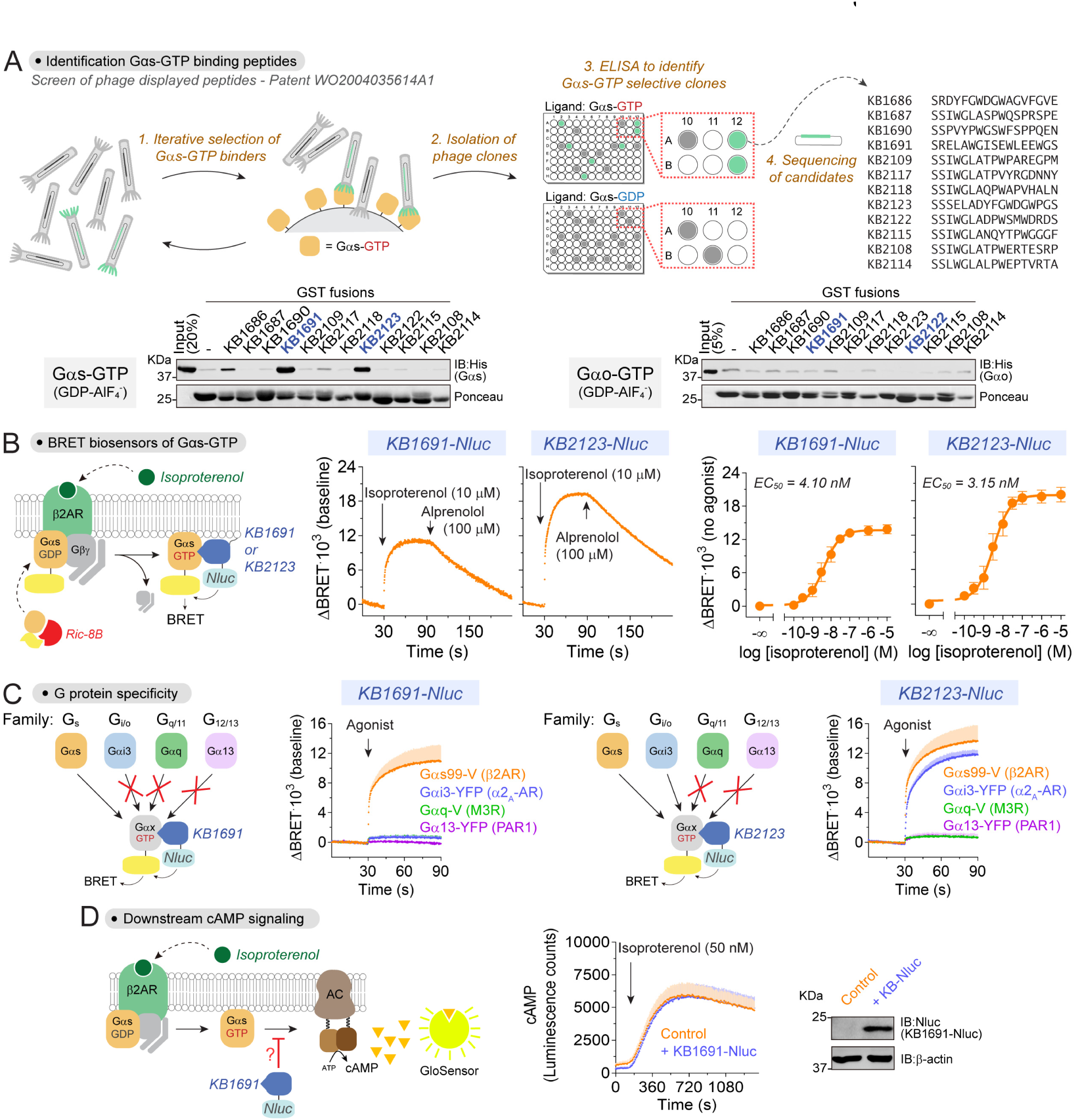
Direct detection of active Gαs in cells with a BRET biosensor. **(A)** Identification of Gαs-GTP peptide binders. *Top,* schematic of phage display screen. *Bottom*, purified GST-fused peptides immobilized on glutathione-agarose beads were incubated with either active, GDP-AlF_4_^-^-loaded Gαs or Gαo, and bead-bound proteins detected by Ponceau S staining or immunoblotting. Results are representative of n ≥ 2 experiments. **(B)** Gαs-GTP biosensor using KB1691 and KB2123 peptides. *Left*, diagram of BRET-based detection of GPCR-mediated activation of Gαs. *Center and left*, BRET was measured in HEK293T cells expressing β2AR, Gαs-99V, Gβγ and either KB1691-Nluc or KB2123-Nluc. A representative kinetic trace from 4 independent experiments in shown in the center panels, and the mean ± S.E.M. of n=4 is shown for the dose-dependence curves on the right. **(C)** Sensors based on KB1691, but not on KB2123, specifically detect Gαs. BRET was measured in HEK293T cells expressing the indicated G protein / cognate GPCR pairs and either KB1691-Nluc or KB2123-Nluc. Cells were stimulated with either 10 μM isoproterenol (for Gαs), 5 μM brimonidine (for Gαi3), 100 μM carbachol (for Gαq), or 30 μM TRAP-6 (for Gα13). Results are mean ± S.E.M. of n=3-4. **(D)** KB1691 does not interfere with Gαs-mediated cAMP production. Luminescence was measured in HEK293T cells expressing the cAMP probe Glosensor and exactly the same components as in panel B, except that KB1691-Nluc was omitted in the control. Results are the mean ± S.E.M. of n=4. A representative immunoblotting result confirming expression of KB1691-Nluc is shown on the right.

### Specific detection of active Gαs but no other G protein type

Next, we asked if the newly identified biosensors were specific for Gαs. To test this, we replaced Gαs-99V with YFP-tagged Gα subunits representative of the other three G protein families (i.e., Gαi3 for G_i/o_, Gαq for G_q/11_, and Gα13 for G_12/13_), and measured BRET responses upon stimulation of cognate GPCRs (**Fig. 1C**). Activation of Gαi3, Gαq and Gα13 under these conditions was confirmed in parallel experiments with previously validated biosensors for each one of these G proteins^30^ (**Fig. S3**). We found that KB1691-based sensors only detected activation of Gαs, whereas KB2123-based sensors detected activation of both Gαs and Gαi3 (**Fig. 1C**). These results demonstrate that biosensors based on the KB1691 sequence, but not on the KB2123 sequence, specifically detect activation of Gαs. For this reason, we focused our subsequent efforts on biosensors based on KB1691.

### Gαs-GTP biosensor does not interfere with downstream signaling

A potential concern of any biosensor is the potential interference with the process it measures. Because Nluc is a bright luciferase that can be expressed at low levels to obtain measurable luminescence signals^41^, we reasoned that expression of the biosensor with Nluc-fused KB1691 would not interfere significantly with G_s_ signaling in cells. To test this, we measured cAMP production upon β2AR stimulation in HEK293T under conditions identical to those used to detect Gαs activation with this sensor, except that a probe for cAMP (GloSensor-22F, Promega)^42^ was co-expressed. We found that the cAMP response detected was not affected by the expression of the Gαs BRET sensor (**Fig. 1D**), indicating no overt effect on GPCR-stimulated G_s_ signaling to downstream effectors.

### Gαs-GTP biosensor reveals specific activation properties of oncogenic Gαs mutants

Having established a biosensor for Gαs-GTP, we set out to leverage it in functional studies, starting with characterizing the activation properties of cancer-associated G protein mutants. GNAS, the gene encoding Gαs, is frequently mutated in cancer at two hotspots corresponding to R201 and Q227 in the protein^43,44^. These mutations lead to Gαs forms that are constitutively active in cells— i.e., they elevate cAMP levels under unstimulated conditions^45^. Both residues are conserved in the nucleotide binding pocket of Gα subunits and participate in GTP hydrolysis (**Fig. 2A**), suggesting that the mechanism by which their mutation leads to constitutive signaling is by locking them in a GTP-bound state. However, *in vitro* results with purified proteins suggest that Q227 mutants are likely to be completely occupied by GTP, whereas R201 mutants might be only partially occupied with GTP^46,47^. Whether this is the case in cells is not known, so we leveraged the newly developed Gαs-GTP biosensor to characterize the activation properties of Gαs R201C and Gαs Q227L. We found that, under unstimulated conditions, Gαs R201C gave a higher BRET signal than Gαs WT, yet it was much lower than the BRET signal for Gαs Q227L (**Fig. 2A**). Moreover, isoproterenol stimulation did not lead to an increase of the BRET signal for Gαs Q227L (**Fig. 2A**), consistent with the interpretation that this mutant is completely occupied with GTP in cells and cannot be further activated. In contrast, isoproterenol stimulation led to an increase of BRET for Gαs R201C, which was not reverted upon an addition of antagonist that fully reverted the isoproterenol response with Gαs WT (**Fig. 2A**). While the latter indicates that Gαs R201C is indeed GTPase-deficient, the former suggests that Gαs R201C is susceptible to further GTP loading upon acute receptor stimulation. Taken together, these results indicate that, as opposed to Gαs Q227L, Gαs R201C is only partially occupied with GTP in cells.

**Figure 2.**
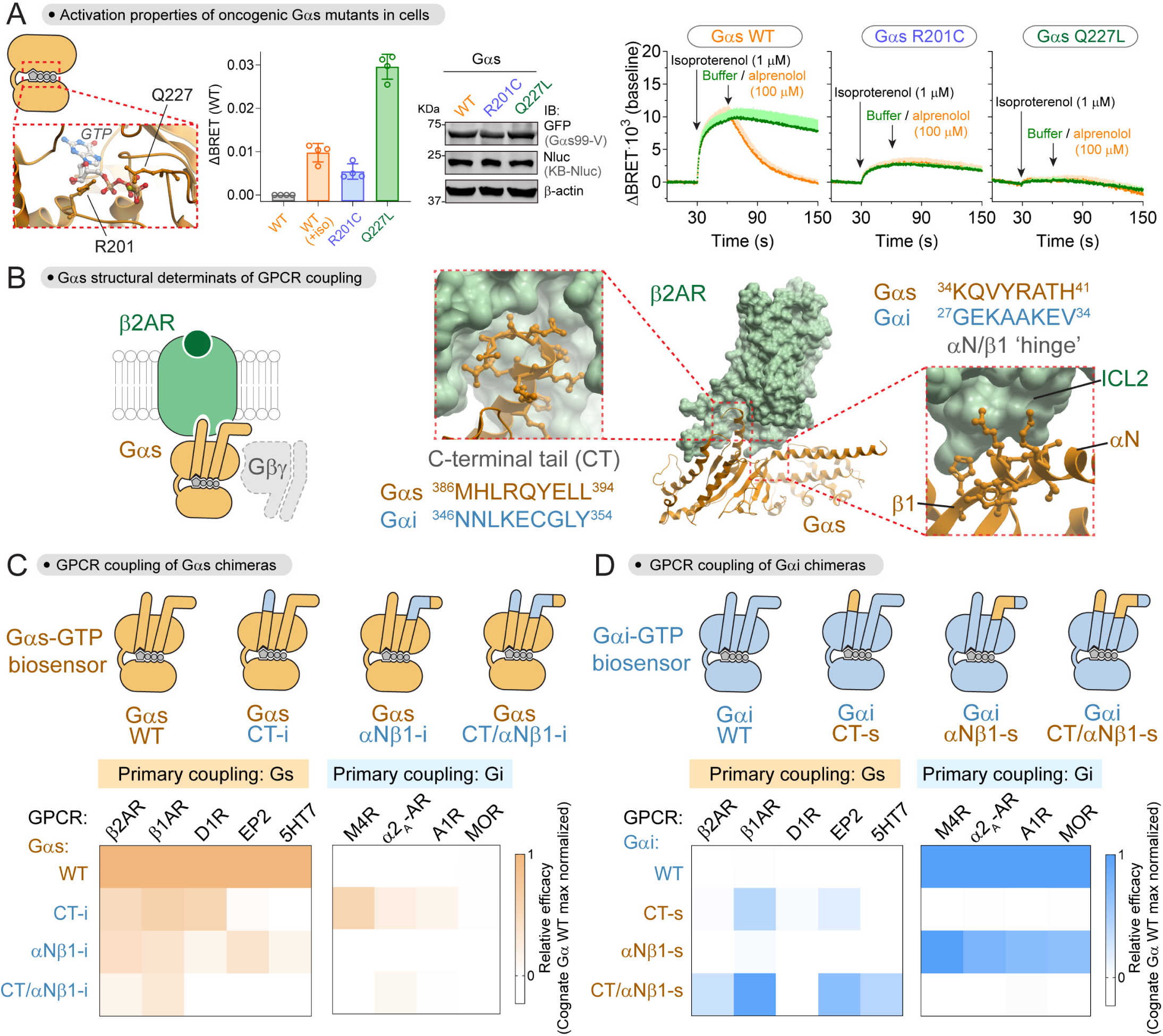
Mechanism of Gαs activation by oncogenic mutations and of G_s_ coupling to GPCRs. **(A)** Activation properties of oncogenic Gαs mutants. *Left,* view of Gαs nucleotide-binding pocket with residues mutated in cancers (PDB: 1AZT). *Right*, BRET was measured in HEK293T cells expressing the same components as in Fig. 1B with KB1691-Nluc, except that Gαs-99V (WT) was replaced as indicated by Gαs-99V R201C or Q227L mutants. Bar graph represents BRET signal in unstimulated cells or cells stimulated with 1 μM isoproterenol relative to unstimulated Gαs-99V WT. Results are the mean ± S.E.M., n=4. Kinetic traces of BRET measurements with cells expressing the same components and treated as indicated. Results are expressed as the difference in BRET from their corresponding unstimulated baselines. Results are the mean ± S.E.M., n=3. The immunoblot is a representative result confirming equal expression of Gαs and KB1691-Nluc. **(B)** View of structural elements of Gαs involved in coupling to β2AR based on their complex structure (PDB: 3SN6). **(C, D)** Contribution of the C-terminal tail (CT) and the αN/β1 ‘hinge’ of Gα subunits to their coupling to GPCRs. BRET was measured in HEK293T cells expressing the indicated YFP-tagged Gα chimeras, GPCRs, and Gβγ with either a Gαs-GTP biosensor (KB1691-Nluc, **C**) or a Gαi-GTP biosensor (KB1753-Nluc, **D**). Heat maps correspond to the efficacy of the BRET responses detected relative to the maximal response observed with the cognate WT G protein. Results are the mean of n=3-4. Full dose-dependence curves are presented in **Fig. S4**).

### Gαs-GTP biosensor reveals G_s_ structural determinants for productive GPCR coupling

Given that the newly developed G_s_ activity biosensor measures the direct product of the catalytic GEF activity of GPCRs, i.e., Gαs-GTP, we set out to leverage it to characterize structural determinants of Gαs required for receptor-mediated activation. While it is well-established that the C-terminal tail of Gα is critical for GPCR coupling, it has become clear that this is not the sole determinant required for efficient receptor coupling^48–51^. Based on the structure of the β2AR-Gs complex and other related GPCR-G protein structures, the αN/β1 hinge region of Gαs makes extensive contact with the receptor and may serve as an allosteric conduit between the receptors and the nucleotide binding pocket of G proteins^52,53^ (**Fig. 2B**). To directly determine the relative contribution of the C-terminal tail (CT) and the αN/β1 hinge to the productive coupling of Gαs to GPCRs, we used the newly generated Gαs-GTP biosensor to measure β2AR-mediated activation of Gαs chimeras in which the CT and/or the αN/β1 hinge had been replaced by the corresponding sequences in Gαi3, which is not activated by this receptor (**Fig. 2C, Fig S4**). As expected, replacement of the CT led to a marked decrease in activation (∼50%) compared to wild-type (**Fig. 2C, Fig. S4**). Interestingly, replacement of αN/β1 hinge led to a similar decrease in activation (∼50%), and the effects became additive when both CT and αN/β1 hinge were replaced (∼95% decrease). These results indicate that not only the CT, but also the αN/β1 hinge, is necessary for efficient coupling of Gαs to β2AR (**Fig. 2C, Fig S4**). Moreover, replacement of the αN/β1 hinge region impaired activation by four other GPCRs known to couple primarily to G_s_ (**Fig. 2C, Fig S4**), indicating that the requirement for the αN/β1 hinge region is a general feature of G_s_-GPCR coupling. Similar experiments were carried out with reciprocal Gαi3 chimeras to assess what elements of Gαs are *sufficient* to permit activation. Replacement of the CT of Gαi3 with the corresponding region of Gαs led to some activation upon stimulation of the β1AR and EP2 (**Fig. 2D, Fig S4**). When both the CT and αN/β1 hinge regions were replaced simultaneously, activation was enhanced for these two receptors compared to the CT chimera, and responses became detectable for two additional GPCRs (β2AR and 5HT7) (**Fig. 2D, Fig S4**). These results highlight the importance of the αN/β1 hinge region of Gα for activation by GPCRs that primarily couple to G_s_. However, we found that this is not the case for GPCRs primarily coupled to G_i_ because replacement of the αN/β1 region of Gαi3 by the corresponding region of Gαs had little or no effect on its activation by four different receptors, while replacement of the CT completely ablated the responses (**Fig. 2D, Fig S4**). Moreover, grafting the αN/β1 hinge region of Gαi3 into Gαs was not sufficient to achieve activation of the non-cognate G protein by any of the four G_i_-linked GPCRs, whereas replacement of the CT led to activation by three of them (**Fig. 2C, Fig S4**). These results indicate that the CT, but not αN/β1 hinge, of Gαi has a prominent role in achieving productive coupling to GPCRs. Taken together, our observations indicate that the αN/β1 hinge region is a structural determinant specifically important for Gαs productive coupling to cognate GPCRs, and that this feature might not be shared by other G proteins.

### ONE vector G protein Optical (ONE-GO) biosensor designs display improved features

Next, we set out to condense all the components of the Gαs-GTP biosensor into a single vector. We reasoned that delivery of the biosensor as a single payload would provide several advantages. For example, it would facilitate implementation without the need for tedious optimization of expression conditions in high-throughput formats or when investigating endogenous GPCRs in poorly transfectable primary cells. Another advantage of delivering the biosensor components as a single payload is that every cell receiving the construct would express all components required for the BRET biosensor^54,55^, thereby increasing the magnitude of the responses detected while simultaneously reducing the total amount of exogenous G protein required (**Fig. 3A**). As an initial step, we reduced the number of components required for detection of Gαs from five to three by removing exogenous expression of Gβ and Gγ, which do not directly participate in the BRET interaction between Gαs-99V and Nluc-fused KB1691 (**Fig. 3A, Fig. S5**). We found that expression of exogenous Gβ and Gγ was dispensable to obtain robust BRET responses with two different GPCRs (**Fig. S5**). This not only indicates that we can reduce the number of components expressed without sacrificing quality of performance, but also that our biosensor detects responses of G protein heterotrimers assembled with endogenous Gβγ.

**Figure 3.**
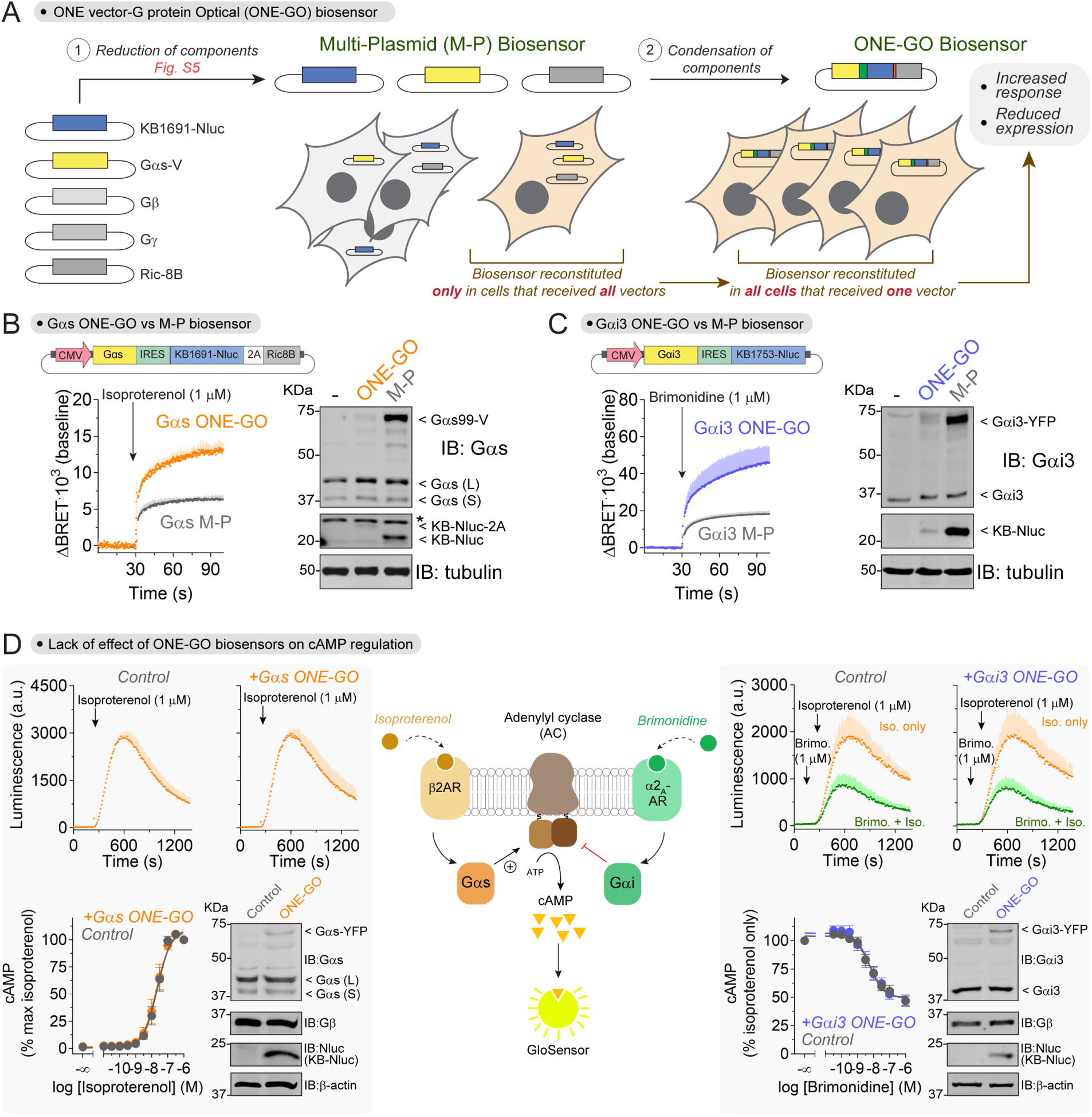
ONE vector G protein Optical (ONE-GO) biosensor designs display improved features. **(A)** Schematic of the process to develop ONE-GO biosensors. **(B-C)** Gαs and Gαi3 ONE-GO biosensor designs provide increased responses and reduced component expression. BRET was measured in HEK293T cells transfected with the indicated single-plasmid ONE-GO biosensors or their multi-plasmid (M-P) counterparts. β2AR or α2_A_-AR were co-expressed for Gαs (**B**) or Gαi3 (**C**), respectively. Results are the mean ± S.E.M. of n=3. A representative immunoblotting result comparing expression of sensor components is shown alongside the BRET traces. **(D)** ONE-GO biosensors do not interfere with GPCR-G protein signaling. Luminescence was measured in HEK293T cells expressing the cAMP probe Glosensor with or without the Gαs ONE-GO sensor (*Left*) or Gαi3 ONE-GO sensor (*Right*). Cells used on the right co-expressed exogenous α2_A_-AR, whereas isoproterenol responses were elicited by endogenous β2AR. Results are the mean ± S.E.M. of n=3. A representative immunoblotting result confirming expression of sensors and lack of changes in endogenous G protein subunits is shown for both ONE-GO sensors.

We then set out to design and optimize a tricistronic Gα-GTP biosensor construct coding for Gαs-99V, KB1691-Nluc, and Ric-8B (**Fig. 3A**), which we named Gαs <u>ONE</u> vector <u>G</u> protein <u>O</u>ptical biosensor (Gαs ONE-GO biosensor). This construct and its derivatives described below were assembled in a lentiviral plasmid backbone ensuring that the size of the payload remained below the limit for lentiviral packaging. We explored designs with two promoters of different strength, SV40 and CMV, in which Gαs-99V was placed directly downstream of the promoter and KB1691-Nluc downstream of an IRES, followed by a self-cleaving T2A sequence and Ric-8B (**Fig. S6A**). We used a modified IRES with low efficiency (IRES*)^56,57^ to achieve a higher acceptor to donor expression ratio, which is expected to improve the dynamic range of BRET responses. Similar designs were also developed for another G protein (Gαi3) using a cognate Gα-GTP detector module (KB1753 peptide^30,58^), with the exception that no chaperone was included (**Fig. S6A**). When compared side by side with the multi-plasmid (M-P) approach, ONE-GO biosensors led to larger responses upon GPCR stimulation (**Fig. 3B, C**, **Fig. 6A**), especially for the designs incorporating the CMV promoter. Moreover, this improvement in performance was accompanied with much lower levels of expression of G protein and Nluc-fused detector modules with ONE-GO compared to M-P (**Fig. 3B, C**). In particular, G protein expression was barely detectable by immunoblotting, and lower than expression of the corresponding endogenous G protein. Overall, these results indicate that ONE-GO biosensors detect robust responses under near-endogenous G protein expression conditions, in which low amounts of exogenous Gα subunits assemble into functional heterotrimers with endogenous Gβγ dimers.

### ONE-GO biosensors do not interfere with GPCR-G protein signaling

To mitigate potential concerns of interference of ONE-GO biosensors with GPCR signaling, we tested their effect on a signaling readout downstream of G protein activation (i.e., cAMP). We found that neither Gαs ONE-GO nor Gαi3 ONE-GO changed the efficacy or potency of GPCR-mediated cAMP responses (**Fig. 3D**). Gαs ONE-GO biosensor expression did not alter isoproterenol-induced cAMP production, whereas Gαi3 ONE-GO did not affect either isoproterenol cAMP production or its inhibition upon stimulation of the α2_A_ adrenergic receptor (α2_A_-AR) (**Fig. 3D**).

### ONE-GO biosensors report activation across G protein families and for many GPCRs

Next, we extended the ONE-GO biosensor design to allow the detection of different Gα-GTP species across all four G protein families (G_s_, G_i/o_, G_q/11_ and G_12/13_) (**Fig. 4A**). We implemented the single plasmid design described above without co-expression of a chaperone by pairing Gα subunits tagged with YFP at the αb-αc loop of the all-helical domain, which tolerates insertions without affecting G protein function^25,30,59^, with Nluc-fused detector modules^30^. We did not attempt to develop constructs for Gαt1, Gαt2, Gαgust, Gαolf, or Gα15 because we lacked validated or suitable detector modules. We generated ONE-GO constructs for Gα11 and Gα14 using GRK2^RH^ as the detector module based on prior evidence^60^, but were not successful in detecting responses with them upon GPCR stimulation. In total, we validated ONE-GO biosensors for 10 different Gα subunits by measuring dose-dependent responses upon stimulation of cognate GPCRs (**Fig. 4A**). These included representative members of all four G protein families, in some cases with diverse members from the same family (e.g., Gαi1 vs Gαz) (**Fig. 4A**).

**Figure 4.**
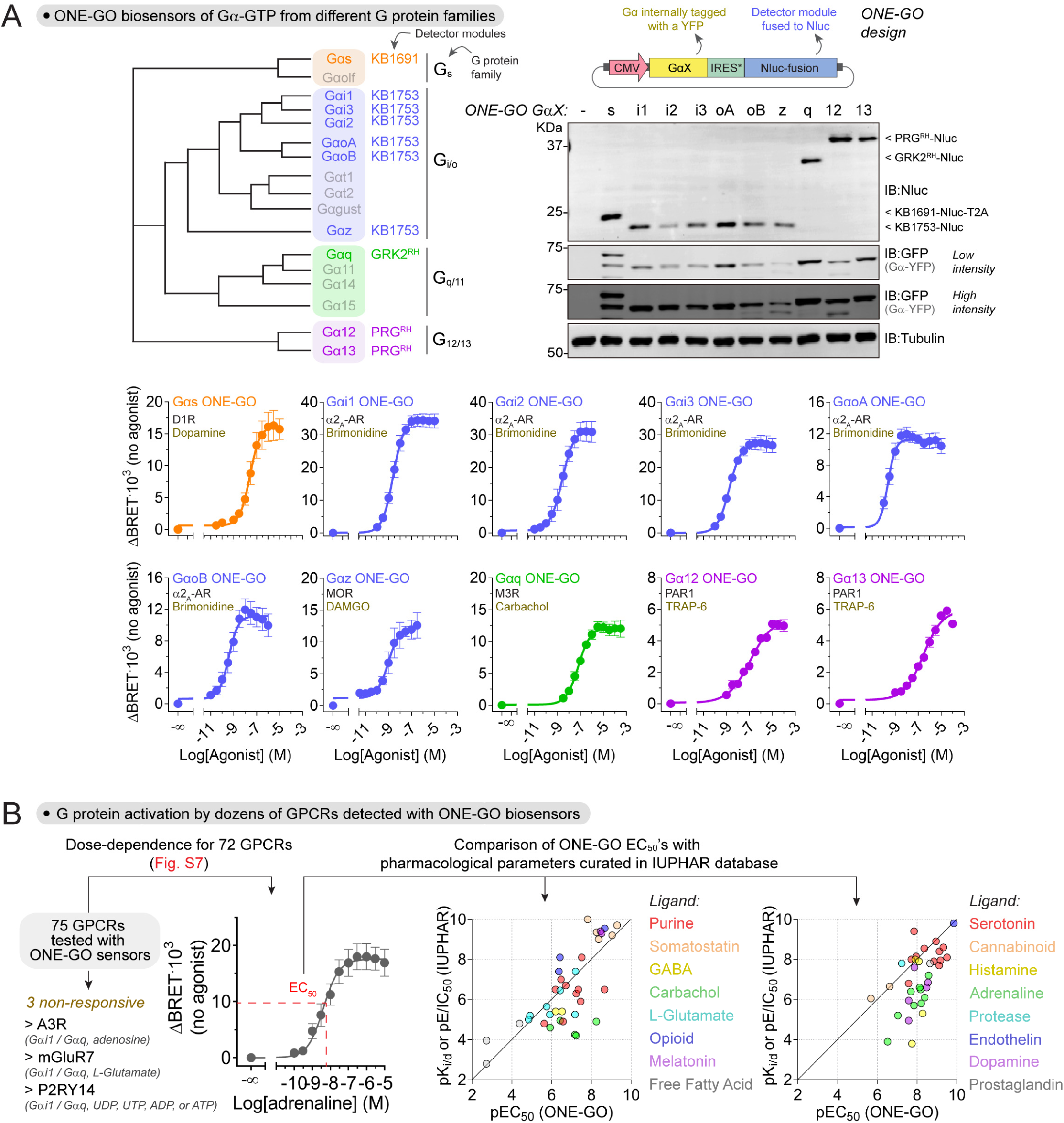
ONE-GO biosensors report activation across G protein families and for many GPCRs. **(A)** Ten ONE-GO biosensors report activity across all G protein families. *Top*, dendrogram of Gα subunits with their corresponding detector modules used in ONE-GO designs, and expression of biosensor components in HEK293T cells. Results are representative of n ≥ 3 experiments. *Bottom*, BRET was measured in HEK293T cells expressing the indicated ONE-GO biosensors and GPCRs. Results are the mean ± S.E.M. of n=3-5. **(B)** ONE-GO biosensors report the activity of dozens of GPCRs. Dose-dependence BRET responses were measured in HEK293T cells for 75 GPCRs to determine EC_50_ values. Results for one representative GPCR (β2AR) are shown (all curves are presented in **Fig. S7**). EC_50_ values were plotted against curated pharmacological parameters (pK_d_, pK_i_, or pE/IC_50_) available in the IUPHAR database.

To further assess the broad utility of ONE-GO biosensors, we implemented them to measure the activity with a large panel of GPCRs and compared results with curated pharmacological parameters (e.g., K_i_ values) available through the International Union of Basic and Clinical Pharmacology (IUPHAR^61^) (**Fig. 4B**). In total, we tested 75 GPCRs with a cognate ONE-GO biosensor (i.e., Gαs, Gαi1, Gαq, or Gα13 ONE-GO), and obtained dose-dependent response curves for >95% of them (72 out of 75 receptors) (**Fig. 4B**, **Fig. S7**). Overall, there was a good correlation between EC_50_ values determined in our assays with ONE-GO biosensors and IUPHAR’s data (**Fig. 4B**). Although comparing our EC_50_’s to the parameters from IUPHAR is limited by the contribution of differences in experimental conditions, these results indicate that detection of GPCR activation by ONE-GO biosensors is broadly reliable and sensitive.

Given the success with Gα-GTP ONE-GO biosensors, we also generated ONE-GO biosensors for the other active signaling species generated upon GPCR stimulation, free Gβγ (**Fig. S6B**). For this, we leveraged the individual components of a previously described Gβγ BRET biosensor^26,27^, which were assembled into a single plasmid such that expression levels of Gα and Gβγ would be similar to favor the formation of functional heterotrimers (**Fig. S6B**). As a proof of principle, we showed that this design detected responses upon stimulation of G_s_ or G_i3_ heterotrimers by a cognate GPCR (**Fig. S6B**).

In summary, our results demonstrate that the ONE-GO biosensor design is broadly applicable to detect GPCR-mediated activation across and within G protein families with high sensitivity.

### Parallel profiling of atypical antipsychotics across a large set of receptors

Given the advantages of delivery as a single genetic cargo and improved dynamic range of ONE-GO biosensors, we set out to demonstrate their suitability for applications in two broad areas: (1) scaling up throughput in experiments with cell lines, and (2) allowing the interrogation of endogenous GPCR activity, even in difficult to transfect primary cells. To start addressing the implementation of ONE-GO biosensors in the first area, we focused our attention on three widely prescribed atypical antipsychotics: brexipiprazole, iloperidone, and cariprazine^62^. Although the exact mechanism of action of these drugs is unknown, the therapeutic benefit of atypical antipsychotics arises from their GPCR polypharmacology, i.e., engaging simultaneously multiple GPCRs, typically with partial agonism/antagonism. Nevertheless, the complex pharmacological profiles of atypical antipsychotics are also believed to underlie the wide range of side effects that vary from one specific drug to another. Although brexipiprazole^63^, iloperidone^64,65^, and cariprazine^66^ have been systematically characterized for binding to large panels of GPCRs, their functional effect as agonists and antagonists has been less systematic, mixing assay readouts and cell lines. We leveraged ONE-GO biosensors to expand and unify the functional characterization of the effect of these three drugs against a panel of 45 GPCRs regulated by neurotransmitters (**Fig. 5A**). Each drug was tested at multiple doses in the absence or presence of a canonical agonist for each GPCR to test agonist or antagonist activity, respectively. For each drug, all receptors were interrogated in parallel for a total of >500 simultaneous measurements (**Fig. 5A**). As expected, the three drugs presented agonist and antagonist activities on numerous GPCRs. From a broad perspective, we found that brexipiprazole and cariprazine had similar profiles of agonism and antagonism, whereas iloperidone differed, as supported by principal component analysis (PCA) of the datasets (**Fig. 5A**). In broad strokes, iloperidone acts on similar targets as brexipiprazole and cariprazine, prominently serotonin, dopamine, and adrenergic receptors, but with more potent antagonist activity and poor agonist efficacy. At a more granular level, we recapitulated several known features of these drugs, including their action on 5-HT1a, 5-HT2a or D2-like dopamine receptors^63–66^, and also observed pharmacological activities not described previously. For example, we identified inverse agonism for iloperidone on 5-HT1b or 5-HT1d receptors, or the full agonist activity of brexipiprazole and cariprazine on the 5-HT1b and 5-HT2c receptors, or of cariprazine on the 5-HT1d receptor (**Fig. 5A**). All three drugs also caused a marked antagonism on the α2_C_-AR not previously described for any of them. Overall, these findings demonstrate that the ONE-GO biosensor platform can be leveraged to rapidly illuminate GPCR pharmacological profiles through functional interrogation of hundreds of conditions in parallel.

**Figure 5.**
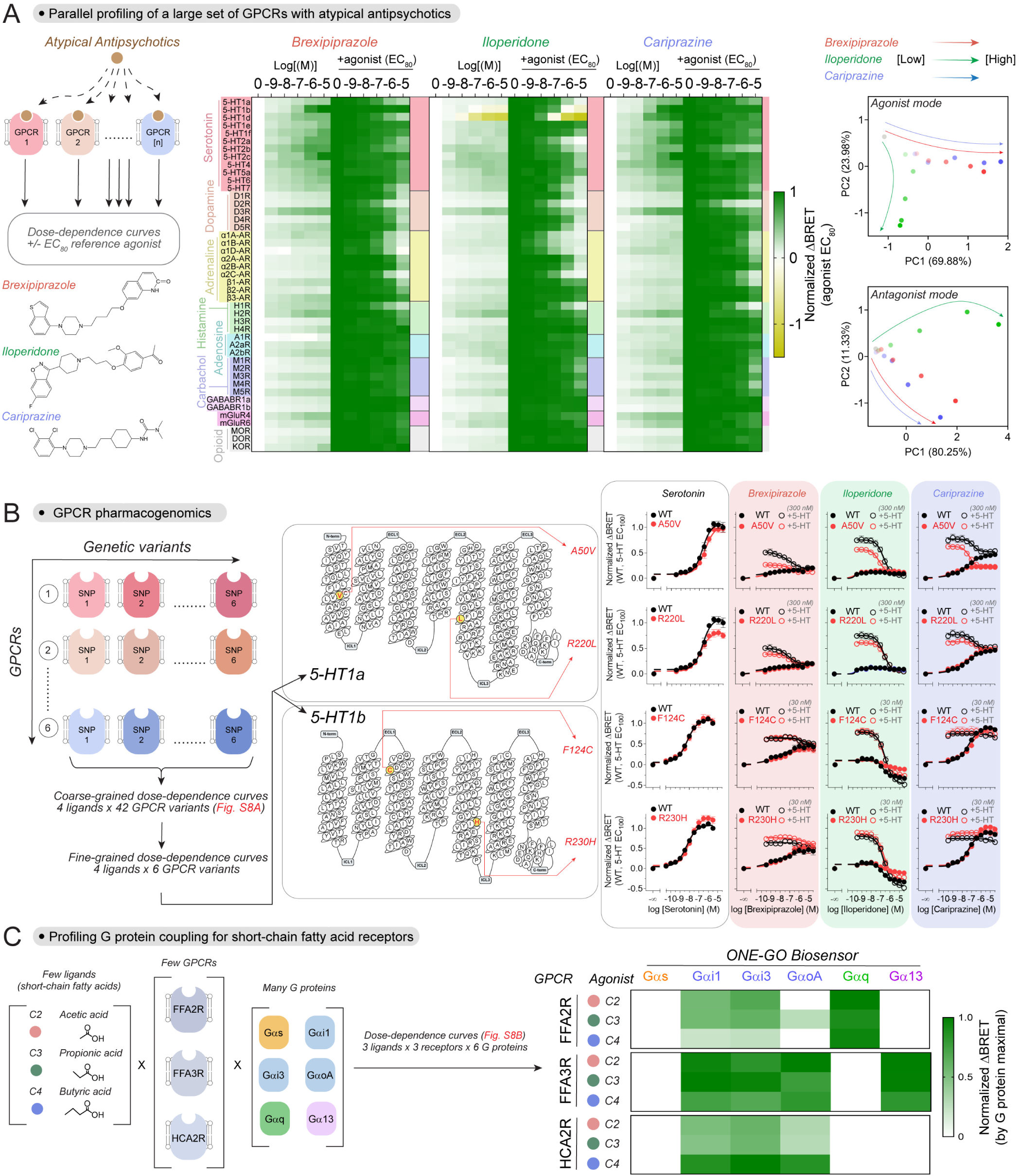
Large-scale parallel interrogation of GPCR activity with ONE-GO biosensors. **(A)** Parallel profiling of atypical antipsychotics across a large set of receptors. *Left*, schematic of the assay and structure of the compounds investigated. *Middle*, BRET responses were measured in HEK293T cells expressing the indicated GPCRs along with a cognate ONE-GO biosensor upon stimulation with the indicated concentrations of brexipiprazole, iloperidone, or cariprazine alone (to measure agonist activity) or in presence of an agonist at its EC_80_ concentration (to measure antagonist activity). Results are the mean of n=3-7. *Right*, principal component analysis (PCA) of the data presented in the heatmaps. See Supplemental Table 1 for agonists and EC_80_ concentrations used. **(B)** Pharmacogenomic profiles of atypical antipsychotics. *Left*, schematic of the assay (coarse-grained curves shown in **Fig. S8A**). *Middle*, Snake plots for 5-HT1a and 5-HT1b showing the location of genetic variations investigated on the right. *Right*, BRET responses were measured in HEK293T cells expressing the indicated GPCRs and the Gαi1 ONE-GO sensor. The indicated compounds were tested by themselves (filled circles) or in presence of 5-HT (open circles). Results are the mean ± S.E.M. of n=3-4. **(C)** Profiling of G protein selectivity across short-chain fatty acid receptors. *Left*, schematic of variables investigated. *Right*, BRET responses were measured in HEK293T cells expressing the indicated GPCR / ONE-GO biosensor combinations upon stimulation with the indicated agonist. Results are maximal responses normalized by biosensor (mean, n=3-6). Full dose-dependence curves in **Fig. S8B.**

### Pharmacogenomics of atypical antipsychotics

Next, we expanded the characterization of atypical antipsychotics with ONE-GO biosensors by assessing how naturally-occurring variants in GPCR sequences influence drug action, a phenomenon known as pharmacogenomics. Pharmacogenomics of GPCR drug targets is believed to account for a significant fraction of adverse side effects or varying drug efficacy across patient populations, but progress in improving prescription precision or clinical trial design has been hampered by the lack of approaches to systematically interrogate the functional consequences of GPCR variants^67–69^. We leveraged ONE-GO biosensors to compare the pharmacological profiles of brexipiprazole, iloperidone and cariprazine on GPCR variants bearing 6 frequent single nucleotide polymorphisms (SNPs) or the most frequent allele (wild-type) for each one of 6 GPCRs modulated by these drugs. We first carried out coarse-grained dose-dependence studies to rapidly identify potential differences by assessing hundreds of experimental conditions in parallel (**Fig. S8A**). Although we recognized that these SNPs could affect total or surface expression of the GPCRs, our goal was to determine if the variant led to different functional outcomes regardless of the underlying cause. Most of the variants led to little or no differences (**Fig. S8A**), whereas four of them were selected for more granular dose-dependence studies to define the putative differences: 5-HT1a A50V, 5-HT1a R220L, 5-HT1b F124C, 5-HT1b R230H (**Fig. 5B**). While the A50V variant had no effect on the efficacy of serotonin, brexipiprazole or iloperidone on 5-HT1a, it made cariprazine a more efficacious antagonist, while reducing its agonist activity (**Fig. 5B**). In contrast, 5-HT1a R220L reduced the efficacy of serotonin, but had no effect on any of the three antipsychotics (**Fig. 5B**). As for, 5-HT1b variants, both F124C and R230H reduced the efficacy of iloperidone as an inverse agonist without changing the effects of brexipiprazole or cariprazine (**Fig. 5B**). R230H but not F124C increased the efficacy of serotonin on 5-HT1b. Overall, these results reveal distinct functional interactions between widely prescribed antipsychotics and naturally-occurring GPCR genetic variants.

### Profiling of G protein selectivity across short-chain fatty acid receptors

Many GPCRs, including those activated by short-chain fatty acids (SCFA), can promiscuously activate different classes of G proteins, which in turn shape cellular responses by modulating different downstream pathways^70,71^. Extracellular SCFA generated by the gut microbiota or as byproducts of eukaryotic metabolism regulate numerous physiological functions via GPCR activation^72–74^. GPCR responses triggered by SCFA are shaped by the intersection of multiple variables, yet the relationship between these variables and the responses has not been systematically deconvoluted using a unified readout^75,76^. The variables consist of multiple ligands [variable 1: acetate, C2; propionate, C3; butyrate, C4] that act on multiple GPCRs [variable 2: FFAR2, FFAR3, and HCAR2] to promiscuously activate G proteins of different families [variable 3] (**Fig. 5C**, **Fig. S8B**). We set out to profile these variables in parallel with ONE-GO biosensors for representative members from each of the 4 G protein families (Gαs, Gαi1, Gαq, Gα13), as well as for different members of the G_i/o_ family (Gαi1, Gαi3, Gαo_A_). None of the receptor-ligand combinations led to a response with Gαs ONE-GO, whereas Gαq activation was only detected for FFAR2 and Gα13 activation was only detected with FFAR3 (**Fig. 5C**, **Fig S8**). Both FFAR2-Gαq and FFAR3-Gα13 responses were similar regardless of the SCFA ligand used. While all three receptors elicited activation of G_i/o_ proteins, they did so with different profiles. First, FFAR2 was a poor activator of Gαo_A_ compared to FFAR3 and HCAR2. The SCFA ligand used also led to different responses across receptors. For example, efficacy and potency of Gαi/o responses decreased with the chain length of the SCFA for FFAR2, whereas HCAR2 showed the opposite pattern (**Fig. 5C**, **Fig S8**). For the Gαi/o responses with FFAR3, the efficacy was similar for the three ligands, but potency was increased for ligands with longer chains (**Fig. 5C**, **Fig S8**). The fact that FFAR2 and FFAR3 show ligand-dependent differences in G_i/o_ responses but not in Gαq or Gα13 responses, respectively, suggests that these receptors may display intrinsically biased G protein responses. In summary, these results provide novel insights into the complex G protein responses elicited by SCFA depending on the combination of available receptors and relative abundance of ligands.

### Time-resolved parallel interrogation of GPCR responses

As another example of potential uses of ONE-GO biosensors to scale up assay throughput in cell lines, we tested the suitability of the system to record multiple kinetic responses to GPCR ligands in parallel, instead of the end-point measurements described above. Time-resolved measurements can be useful not only to evaluate changes in activation/deactivation kinetics, but also to provide multiple modes of pharmacology like agonism, antagonism, and allosteric modulation from a single well^77,78^. We measured time-resolved responses simultaneously across 48 wells with two different receptors (α2_A_-AR and GABA_B_R), obtaining high reproducibility scores (i.e., Z’ > 0.9), and resolving an expected delay in G protein deactivation kinetics after disabling RGS GAP mediated regulation with mutation G183S in Gαi3^79,80^ (**Fig. S8C**).

### Detection of endogenous GPCR activity in cells lines

Next, we shifted our attention to the detection of responses elicited by endogenous GPCRs. As a first test, we generated stable HeLa cells expressing Gαi3 ONE-GO or Gαi3/Gβγ ONE-GO biosensors by lentiviral transduction followed by fluorescence-activated cell sorting (**Fig. S9B, C**). We previously showed that activation of endogenous α2 adrenergic receptors in these cells led to detectable Gαi-GTP or free Gβγ responses with a different biosensor design named BERKY^30^. We found that stimulation with an α2-adrenergic agonist gave rise to robust dose dependent responses in cells expressing different levels of Gαi3 ONE-GO or Gαi3/Gβγ ONE-GO biosensors, and that these responses were much larger than those observed previously in the same cells with Gαi*-BERKY3 or Gβγ-BERKY3 biosensors, respectively^30^ (**Fig. S9B, C**). Similarly, we transduced SH-SY5Y cells with a lentivirus for the expression of the Gαi3 ONE-GO biosensor and compared responses triggered by endogenously expressed opioid receptors with those observed in a stable SH-SY5Y cell line bearing the Gαi*-BERKY3 biosensor^30^ (**Fig. S9C**). We found that the ONE-GO biosensor outperformed the BERKY biosensor by eliciting a response almost 10-fold larger. Overall, these results demonstrate that transient or stable expression of ONE-GO biosensors allows the detection of responses triggered by different GPCRs expressed endogenously in two separate cell lines, and that these responses largely exceed those measured by other previously described biosensors suitable for the detection of endogenous GPCR responses.

### Endogenous GPCR activity across a wide palette of primary cells

While detection of responses elicited by endogenous GPCRs is a desirable feature, cell lines might not always recapitulate physiologically-relevant conditions. For this reason, we evaluated ONE-GO biosensors to detect endogenous GPCR responses in primary cells, which should better retain the distinctive features of well-differentiated cells compared to immortalized or transformed cell lines. First, we succeeded in detecting kinetic and dose-dependent responses triggered by endogenous adenosine or protease activated receptors in primary human cardiac fibroblasts transduced with lentiviruses of ONE-GO biosensors for G proteins representative for each one of the four families (**Fig. 6**). Motivated by this, we expanded our studies to five other primary cell types representing different organs from both human and mouse origin: human bronchial smooth muscle cells, human umbilical vein endothelial cells, mouse lung fibroblasts, mouse astroglial cells, and mouse cortical neurons. To achieve neuron-specific expression, the CMV promoter was replaced by the synapsin I promoter in the lentiviral constructs. **Figure 6** shows examples of responses detected in all these cell types, which account for all G protein families and at least 10 general types of receptors for diverse ligands. **Figure S10** illustrates additional examples of ligand-biosensor pairs, including Gαo_A_ responses, and controls assessing the specificity of the responses observed. For example, Gαi/o responses were blocked by pertussis toxin^81^, and Gαq responses by YM-254890^82^, whereas none of these treatments affected Gα13 responses, as expected (**Fig. S10**). Given the lack of inhibitors for Gαs, the specificity of the responses detected was determined by the use of selective antagonists for the presumed G_s_-coupled GPCRs (**Fig. S10**). These results demonstrate the suitability and broad applicability of ONE-GO biosensors for the detection of endogenous GPCR responses mediated by any general type of G protein across an ample palette of distinct primary cell types.

**Figure 6.**
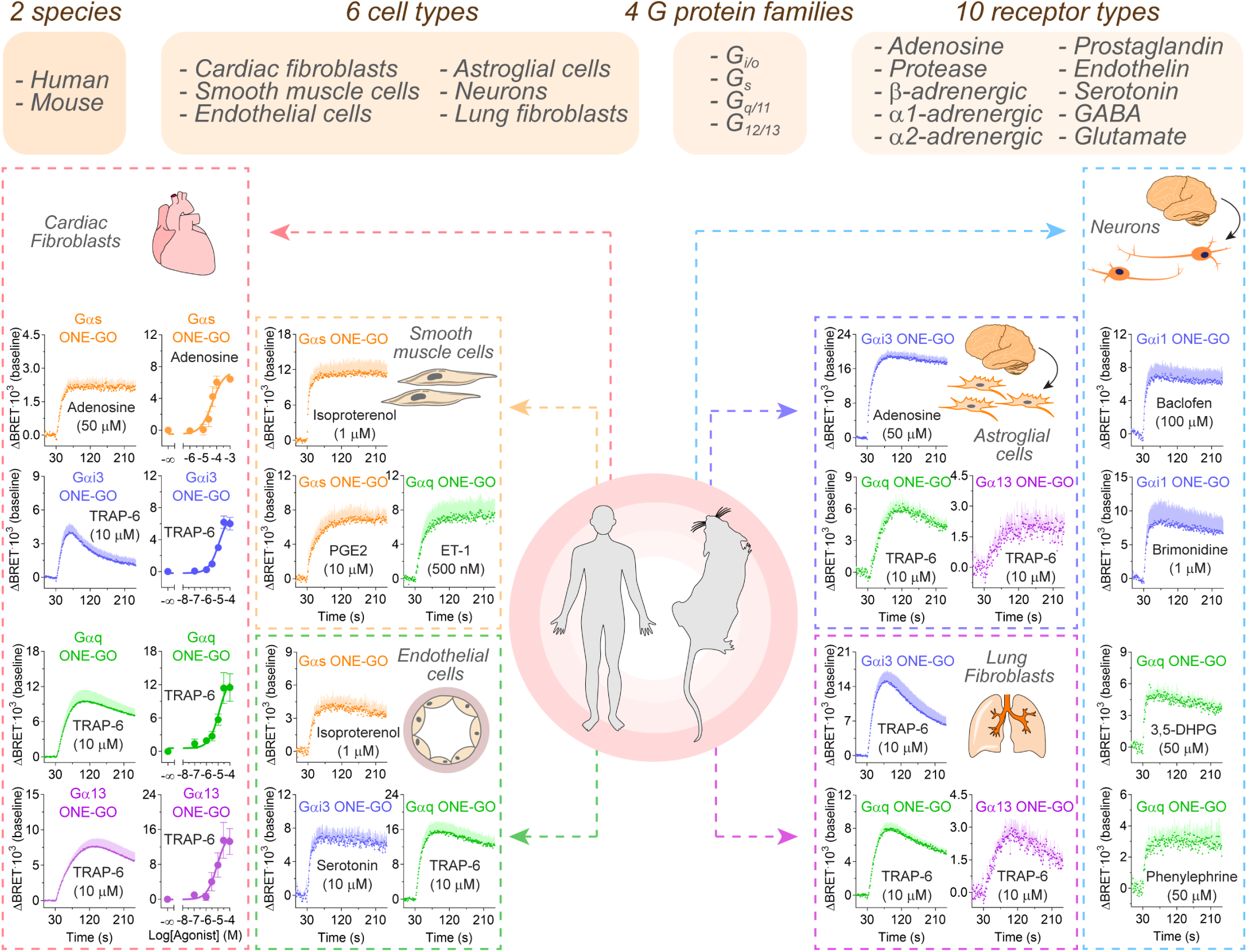
Detection of endogenous GPCR activity across a wide palette of primary cells with ONE-GO biosensors. *Top*, summary of responses triggered by endogenous GPCRs detected with ONE-GO biosensors for all G protein families in multiple human and mouse primary cells of different origins. *Bottom*, BRET responses were measured in the indicated primary cells transduced with ONE-GO biosensors. Results are the mean ± S.E.M. of n=3-6. Additional examples and controls measured in parallel in **Fig. S10**.

### Cell type specific G protein selectivity profiles of protease-activated receptor 1

A poorly understood but critical aspect of GPCR regulation is its context dependence. In other words, how does GPCR behavior depend on the cell type or state in which it is expressed? Since our results above validated that ONE-GO biosensors are suitable to answer this question, we set out to elaborate on it. In the next sections, we provide three examples of how measuring endogenous GPCR activity reveals context-specific behavior; from how a receptor changes its profile of activation of different G protein families depending on cell type, to discrimination of G protein subtypes within the same family for some receptors in neurons, to remodeling of G protein activation profiles upon induction of disease-mimicking cell transformation.

Protease activated receptor 1 (PAR1) is known to promiscuously couple to G proteins of the G_i/o_, G_q/11_ and G_12/13_ families^70,71^, which trigger distinct downstream signaling cascades. Although G protein coupling specificity has been systematically addressed with unified readouts in cell lines expressing exogenous PAR1^70^, this has not been done for endogenously expressed PAR1. We stimulated endothelial cells, lung fibroblasts, cardiac fibroblasts and astroglial cells with a PAR1-specific agonist and measured activation of Gαq, Gαi3, or Gα13 by transducing the corresponding ONE-GO biosensors (**Fig. 7A**). All cell types exhibited a robust Gαq response, but the relative strength of the Gαi3 and/or Gα13 responses varied across cell types giving rise to distinct profiles of G protein activation (**Fig. 7A**, *left*). For example, Gαi3 responses were very weak or absent in endothelial cells and cardiac fibroblasts, whereas Gα13 responses were strong in cardiac fibroblast but only modest in the other three cell types (**Fig. 7A**). The different response profiles could not be explained by differences in relative expression of the three biosensors, as determined by luminescence measurements (**Fig. 7A**, *right graphs*). These results reveal that the profile of PAR1 coupling to G proteins of different families depends on the cell type in which the receptor is expressed.

**Figure 7.**
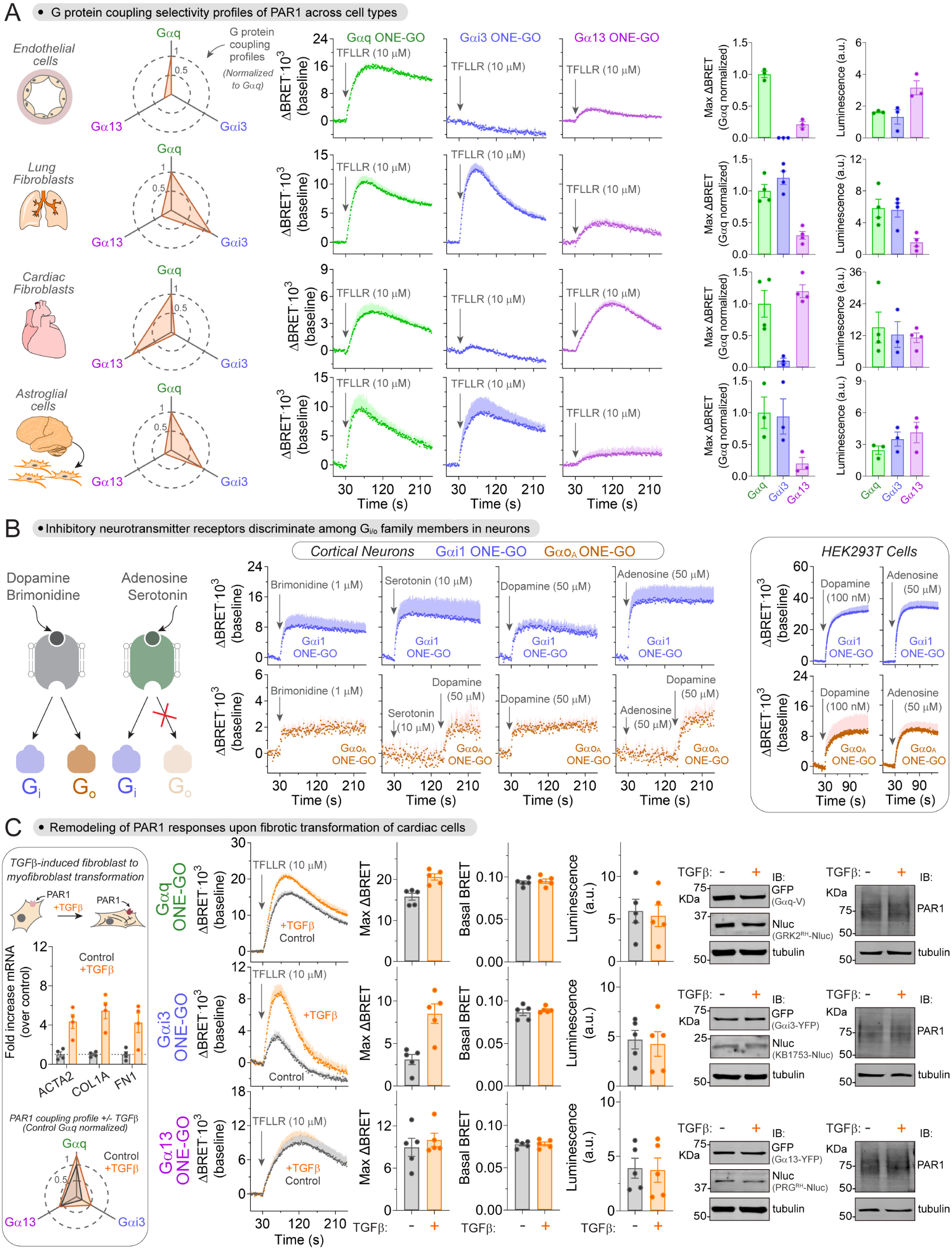
ONE-GO biosensors reveal context-dependent activity of endogenous GPCRs. **(A)** Cell type-dependent G protein selectivity profiles of protease-activated receptor 1. *Left*, spider plot summarizing PAR1 G protein activation profiles for each cell type. *Middle*, BRET responses were measured in each one of the 4 primary cell types transduced with the indicated ONE-GO biosensor. *Right*, maximal BRET responses normalized to Gαq ONE-GO and raw luminescence counts indicative of biosensor expression. Results are the mean ± S.E.M. of n=3-4. **(B)** Discrimination across G_i/o_ isoforms by neuroinhibitory GPCRs in primary neurons. *Left*, BRET responses were measured in primary mouse cortical neurons transduced with Gαi1 ONE-GO (top row, blue) or Gαo_A_ ONE-GO (bottom row, brown) biosensors stimulated with the indicated neurotransmitters. Results are the mean ± S.E.M. of n=3-6. *Right (box)*, BRET responses were measured in HEK293T cells expressing either the D2R or the A1R along with the indicated ONE-GO biosensor. Results are the mean ± S.E.M. of n=3. **(C)** Myofibroblast transformation remodels the G protein selectivity profile of protease activated receptor 1. *Left (box)*, confirmation of TGFβ-induced myofibroblast transformation by RT-qPCR (mean ± S.E.M. of n=4), and spider plot summarizing PAR1 G protein activation profiles before and after myofibroblast transformation (mean, n=5). *Middle*, BRET responses were measured in human cardiac fibroblasts transduced with the indicated ONE-GO biosensors and treated (orange) or not (grey) with TGFβ. Bar graphs represent from left to right: maximal BRET responses normalized to Gαq ONE-GO, BRET signal in unstimulated cells, and raw luminescence counts indicative of biosensor expression. Results are the mean ± S.E.M. of n=5. *Right*, representative immunoblots showing no difference in expression of the biosensor components (Gα, Nluc) or PAR1 upon TGFβ treatment Results are representative of n=3 experiments.

### Some neuroinhibitory GPCRs discriminate across G_i/o_ isoforms in primary neurons

A widely held tenet based on studies of GPCRs exogenously expressed in cell lines is that those that couple to G_i/o_ proteins do not discriminate between members of the same family like Gαi and Gαo isoforms^70^. For example, a systematic analysis of G protein coupling selectivity in HEK293 cells revealed that out of 104 receptors coupling to Gαi isoforms, only 5 did not couple to Gαo isoforms^70^. We investigated Gαi1 and Gαo_A_ activation using ONE-GO biosensors in mouse cortical neurons upon stimulation with concentrations of agonists known to maximally stimulate specific types of G_i/o_ coupled GPCRs, like α2-adrenergic, 5-HT1 serotonin, D2-like dopamine, and A1 adenosine receptors. While we detected robust Gαi1 responses in all conditions, Gαo_A_ responses were detected only upon stimulation with brimonidine (α2 adrenergic agonist) and dopamine (**Fig. 7B**). The lack of Gαo_A_ response for serotonin and adenosine could not be attributed to poor health of the cell preparations because subsequent stimulation with dopamine in the same recordings led to responses (**Fig. 7B**). It is also unlikely that the absence of response is due to lack of sensitivity, given that Gαi1 responses with serotonin and adenosine are even larger than with brimonidine and dopamine (**Fig. 7B**), so one would have expected proportionally larger Gαo_A_ responses with the former pair of receptors compared to the latter. We confirmed that stimulation of A1 adenosine receptors exogenously expressed in HEK293T cells led to activation of both Gαi1 and Gαo_A_, and that the relative activation of Gαo_A_ compared to Gαi1 is equivalent to that observed upon activation of the D2 dopamine receptor (**Fig. 7B**, *right box*), which contrasts with the observations in neurons. We conclude that some G_i/o_ coupled GPCRs endogenously expressed in primary cortical neurons can discriminate between Gαi and Gαo isoforms.

### Myofibroblast transformation remodels the G protein selectivity profile of protease-activated receptor 1

Next, we leveraged the accessibility of investigating endogenous GPCRs with ONE-GO biosensors to characterize changes in signaling behavior in the context of disease. For this, we focused on PAR1 responses in a model of fibrotic transformation of cardiac fibroblasts. Cardiac fibroblasts are the most abundant cell type in the heart and their conversion into myofibroblasts is a hallmark that underlies heart failure upon fibrotic transformation^83,84^. Although PAR1 receptors contribute to fibrotic transformation in the heart and other tissues^85–89^, whether changes in their signaling behavior accompany the process of transformation and the specific features of these changes are unknown. To address this question, we measured activation of Gαq, Gαi3 and Gα13 with ONE-GO biosensors upon stimulation with a PAR1-specific agonist in a widely used model of fibrotic transformation (**Fig. 7C**). This model consists of inducing the conversion of quiescent cardiac fibroblasts into myofibroblasts by TGFβ stimulation^90,91^, which we confirmed in our hands by detecting the upregulated expression of the fibrosis markers α-smooth muscle actin (ACTA2), collagen I (COL1A), and fibronectin (FN1) (**Fig. 7C**, *left box*). Compared to quiescent cardiac fibroblasts, transformed myofibroblasts displayed a modest increase in Gαq activation, a marked increase in Gαi3 activation, and no difference in Gα13 activation (**Fig. 7C**). These differences could not be attributed to differences in expression of the components of the biosensors or the PAR1 receptor, as assessed by immunoblotting and/or measurements of luminescence (**Fig. 7C**). The lack of difference in basal BRET ratios also indicates that fibrotic transformation does not alter G protein activity under unstimulated conditions (**Fig. 7C**). Overall, these results not only indicate that PAR1 signaling is altered upon fibrotic transformation, but that the changes consist of a remodeling of the relative strength of activation of different G protein types, rather than a global upregulation or downregulation of the receptor responses.

## DISCUSSION

The main advances provided by this work are both technical and conceptual. We generated a set of tools to interrogate GPCR activity with high fidelity and ease even when the receptors are expressed endogenously in their native cellular environment and leveraged these tools to provide new insights into GPCR biology and pharmacology. The ONE-GO biosensor constructs generated here and their designs have been made publicly available through Addgene as an open-source platform to facilitate its adoption and modification by others. The virtually universal applicability of ONE-GO biosensors is supported not only by their demonstrated suitability for a broad set of G proteins and receptors, but also by their implementation across a wide range of cell types (primary cultures and immortalized lines), scalable assay formats (endpoint or kinetic), and gene delivery approaches (transient transfection/ transduction or stable genomic integration). An important concept put forth by our findings with ONE-GO biosensors is that GPCR signaling is hardwired in a context-dependent manner, e.g. in specific cell types, even at the level of the signal transduction event most proximal to ligand-mediated receptor activation (e.g., loading of GTP on G proteins). The phenomenon of context-dependent signaling hardwiring, also known as *system bias*^20^, has been a concern in GPCR drug discovery because it may underlie the lack of consistent translation of findings *in vitro* into the desired pharmacological properties *in vivo*^13,20^. For example, ligand-induced signaling bias currently identified for a given drug candidate *in vitro* using mainstay approaches with transfected cell lines might become negated by system bias when the receptor is expressed endogenously in the target tissue. This issue could be alleviated by implementing ONE-GO biosensors to test drug candidates *in vitro* in cellular systems more relevant to the intended final application, like a primary cell type endogenously expressing the GPCR of interest. A similar approach could also open an avenue for personalized medicine by investigating endogenous GPCR activity in cells differentiated from induced pluripotent stem cells of patients. Other conceptual advances reported here by the implementation of new tools include the elucidation of structure-function relationships in G protein activation, and the discovery of unknown features of the pharmacological or pharmacogenomic profiles of widely prescribed drugs (antipsychotics) or of natural ligands (SCFA).

While our findings of context-dependent signaling reveal that system bias is a prevalent feature of endogenous GPCRs, we can only speculate about the possible causes. For example, although there could be differences in expression of PAR1 across the 4 cell types in which it was investigated, this is unlikely to cause the differences observed because these are changes in the *relative strength* of activation of different G proteins. The possible influence of GPCR and G protein stoichiometry was more definitely ruled out in the experiments testing PAR1 G protein activation remodeling upon fibrotic transformation, in which we confirmed no changes in the abundance of the different signaling components. Similarly, changes in receptor and/or G protein levels cannot explain our observations in neurons. We envision at least three general factors that could shape GPCR responses in a context-dependent manner. One is post-translational modifications. For example, it has been reported that glycosylation of the PAR1 receptor differentially affects signaling cascades triggered by distinct G proteins^92^. Another factor is compartmentalization. For example, GPCRs distributed to specific subcellular structures (e.g., synapses in neurons) or within specific signaling complexes might have different accessibility to specific G proteins^25,93–96^. A third factor is the presence of molecules that directly affect the activity of GPCRs or specific G proteins, which can differ across cell types and/or states. For example, GPCRs can be modulated by lipids like cholesterol or phosphatidylinositol 4,5-bisphosphate^97–100^, or by proteins like Receptor Activity-Modifying Proteins (RAMPs)^101–103^, and G proteins can be regulated by GAPs^104^, GDIs^105^, and GEFs^106–109^, among others^110,111^.

An advantage of ONE-GO biosensors is that they only require the expression of low amounts of exogenous Gα subunits, which bind to endogenous Gβγ to assemble functional G protein complexes that can be activated by GPCRs. On one hand, this prevents the loss of fidelity imposed by the use of specific combinations of exogenous Gβ and Gγ subunits, which alters the signaling profiles of GPCRs^29,112^. On the other hand, this minimizes the potential impact of gross overexpression on the signaling system under investigation, as demonstrated by the lack of measurable effects of ONE-GO biosensors on downstream signaling to modulate cAMP. However, it is still a limitation over other existing biosensor systems like BERKY^30^ that an exogenous G protein subunit needs to be expressed. It is therefore important for investigators to tailor the use of a given biosensor platform to their question of interest. In cases in which fidelity of the response is of utmost importance, BERKY biosensors are more suitable due to the complete absence of exogenous G proteins. However, there might be instances in which the larger dynamic range of ONE-GO biosensors compared to BERKY might justify a minimal compromise of fidelity due to expression of low amounts of Gα. Other limitations to keep in mind for ONE-GO biosensors is that the tagging of Gα could influence their behavior, although this is unlikely based on previous observations^25,30,59^, and that there are no ONE-GO biosensors for some Gα subunits. The latter concern is mitigated by the fact that G protein activation profiles described to date for >100 GPCRs^70^ indicate that their ability to activate G proteins would be captured by at least one of the ONE-GO biosensors currently available.

## ACKNOWLEDGEMENTS

This work was primarily supported by NIH grant R01GM147931 (to MG-M), but also by grant R01NS117101 (to MG-M). RJ is supported by a Predoctoral Fellowship from the American Heart Association (898932). AL is supported by a F31 Ruth L. Kirschstein NRSA Predoctoral Fellowship (F31NS115318). JZ was supported by a Dahod International Scholar Award. HZ was supported by the American Heart Association (23CDA1050577). We thank Dr. A. Belkina and the Boston University Chobanian & Avedisian School of Medicine Flow Cytometry Core for access to FACS resources. We thank the following investigators for providing DNA plasmids: N. Lambert, H. Yano, K. Martemyanov, J. Blumer, P. Polgar, J. Levitz, S. Schulz, P. Slessinger and A. Kovoor. We thank N. Lambert for sharing the Gα protein cartoon used in some figures.

## Author Contributions

R.J., M.M., J-C.P., A.L., E.G., J.Z., C.P., and M.G-M. conducted experiments. R.J., M.M., and M.G-M. designed experiments and analyzed data. H.Z., M.D.L., and J.C.W. provided critical reagents and/or protocols. R.J. and M.G-M. wrote the manuscript with input from all authors. M.G-M. conceived and supervised the project.

## Methods

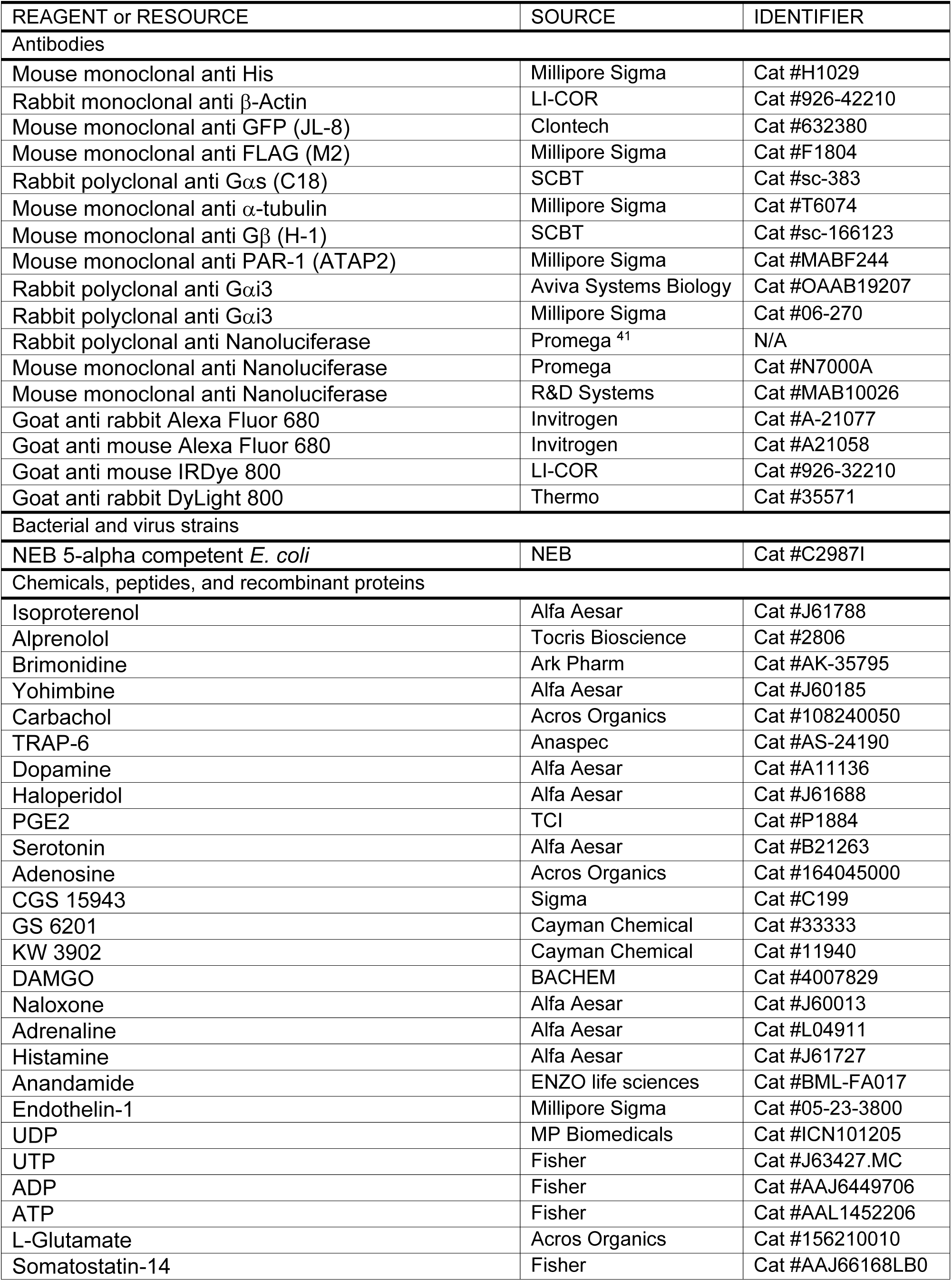
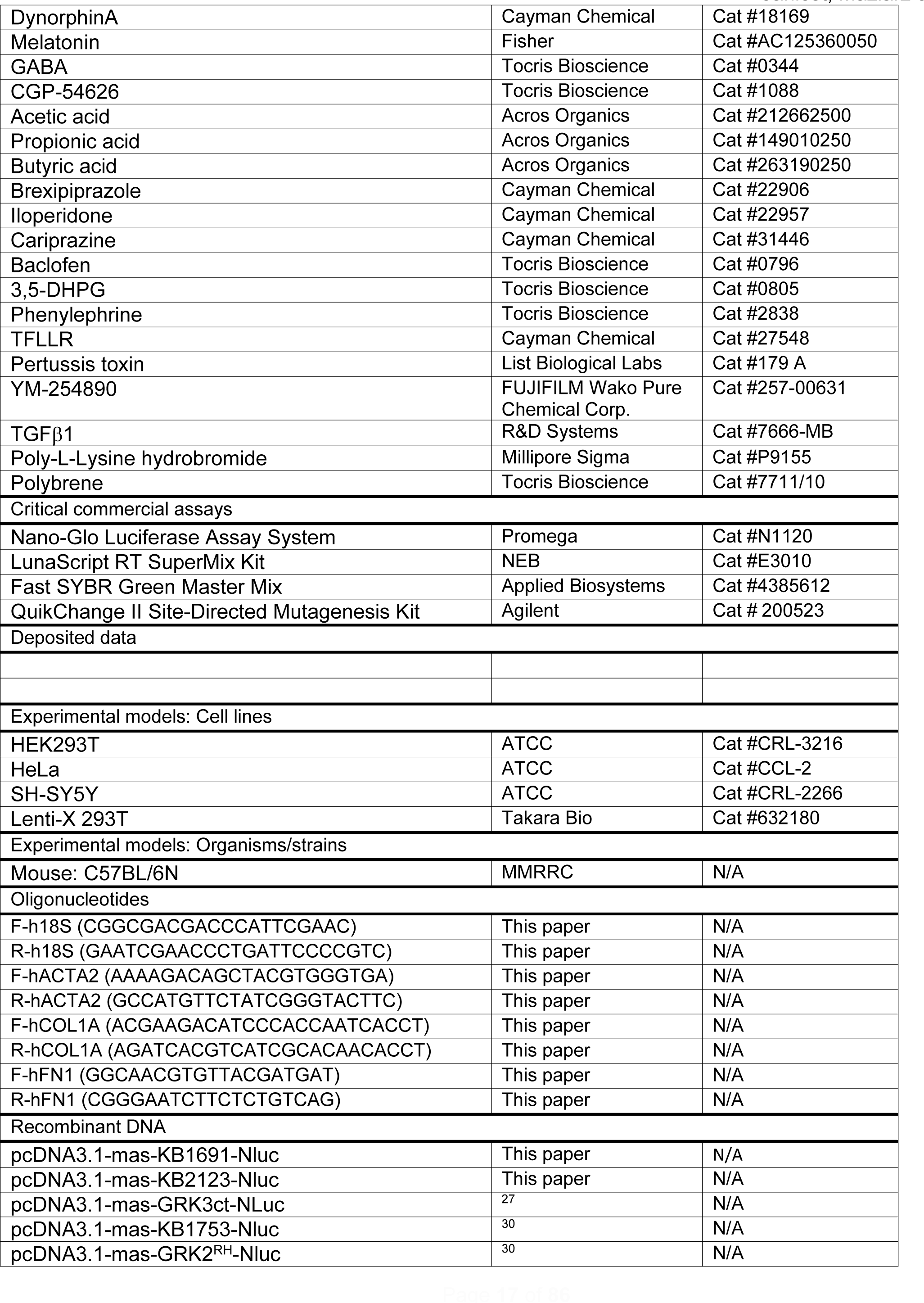
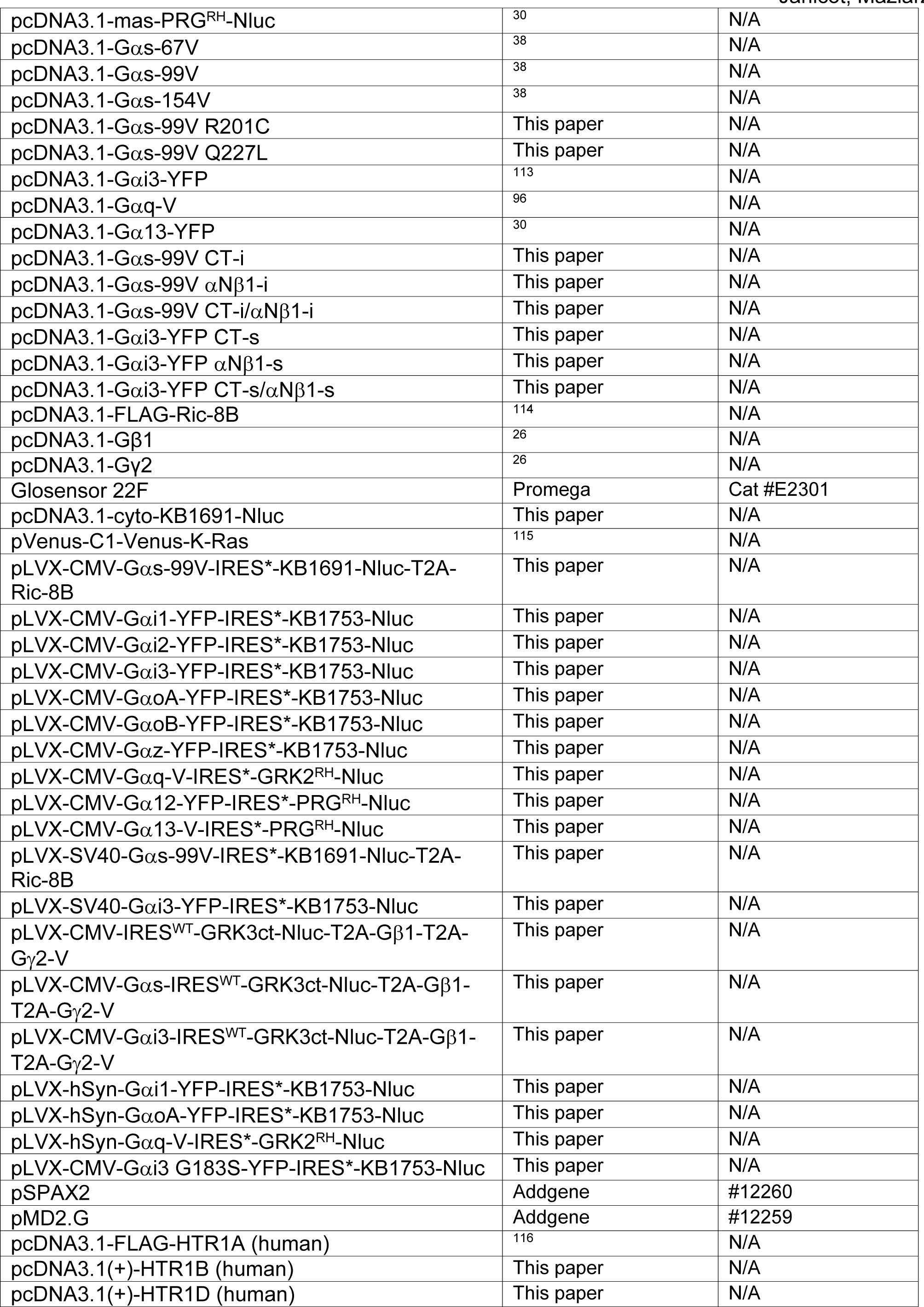
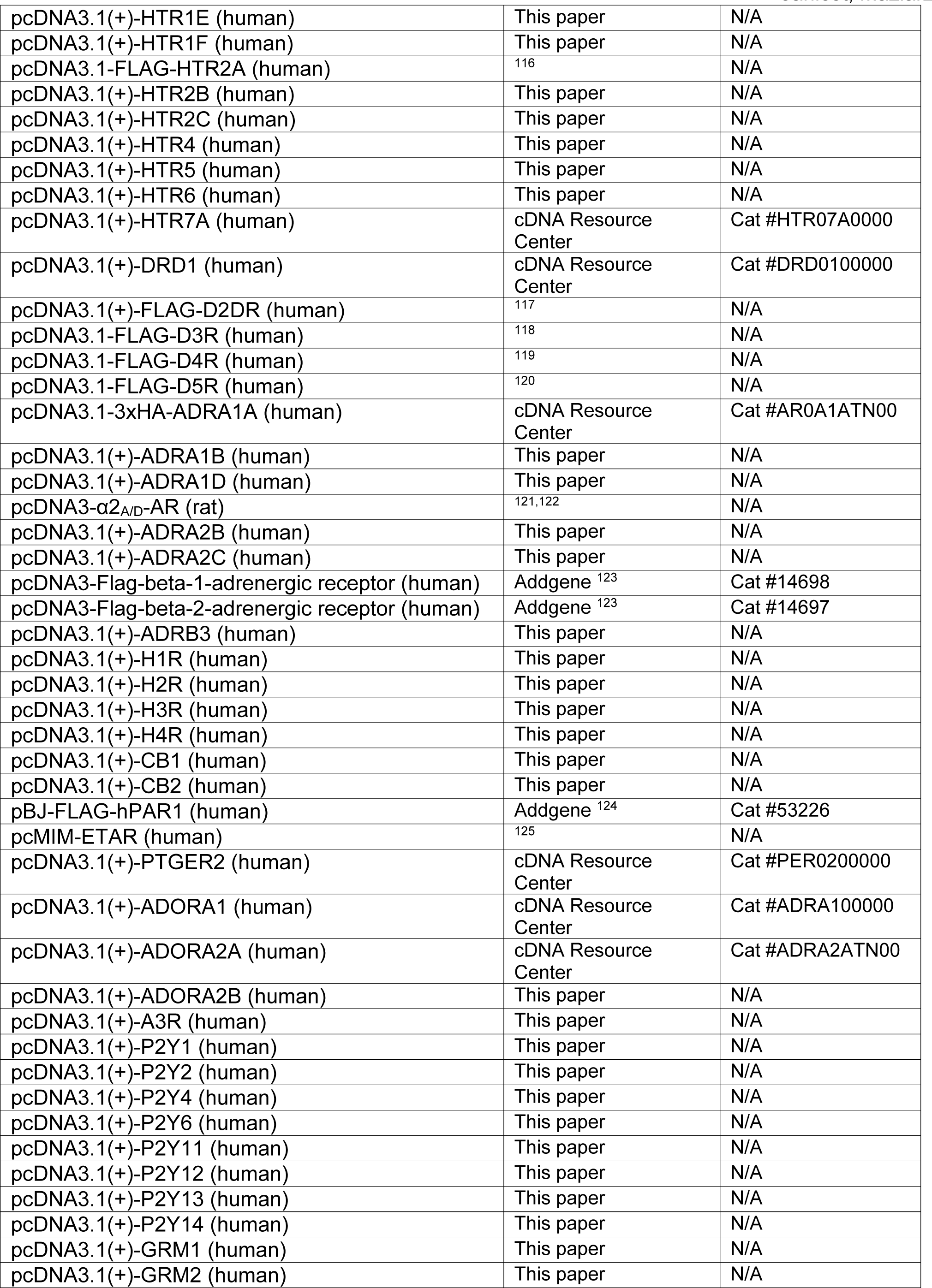
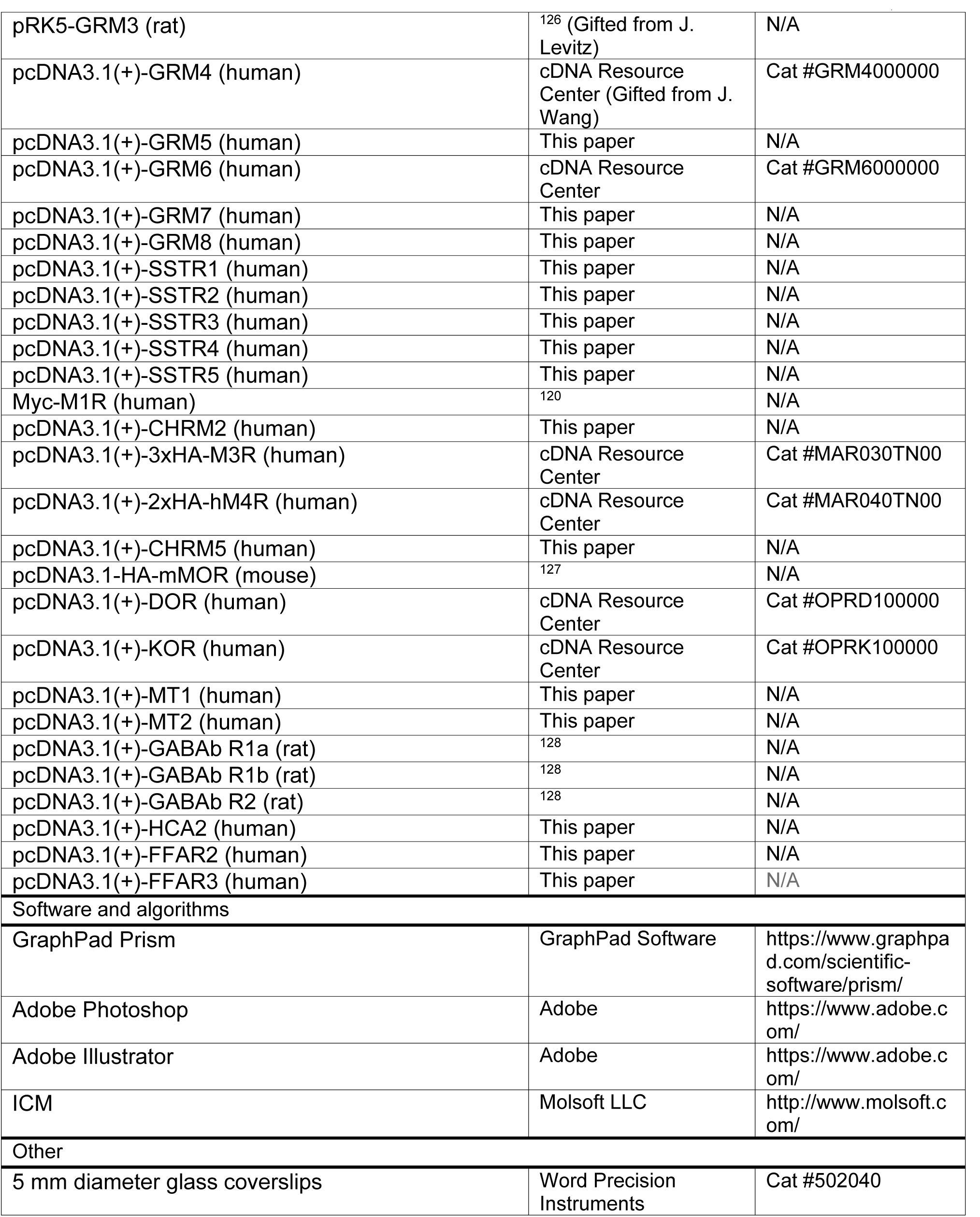

### Experimental Model and Subject Details

#### Cell lines

HEK293T cells (ATCC cat# CRL-3216,) and HeLa cells (ATCC cat# CCL-2) were grown at 37°C, 5% CO_2_ in DMEM (Gibco cat# 11965-092) supplemented with 10% FCS (HyClone cat# SH30073.03), 100 U/ml penicillin, 100 μg/ml streptomycin, and 2 mM L-glutamine. SH-SY5Y cells (ATCC cat# CRL-2266) were grown at 37°C, 5% CO_2_ in DMEM supplemented with 15% heat-inactivated FCS, 100 U/ml penicillin, 100 μg/ml streptomycin, and 2 mM L-glutamine. SH-SY5Y stably expressing Gαi*-BERKY3^30^ were grown in the same medium as naïve cells supplemented with 100 μg/ml hygromycin.

#### Institutional approval for mouse experiments

All animal procedures were approved by the Institutional Animal Care and Use Committee (IACUC) at Boston University Chobanian & Avedisian School of Medicine (PROTO202000018). C57BL/6N wild-type mice were from an in-house colony originally established with animals obtained from the Mutant Mouse Resource & Research Centers (MMRRC) at UC Davis.

#### Mouse primary cortical & striatal neuron culture

Neurons were isolated from neonatal mouse brains as previously described^129^ with modifications. Newborn mouse pups (P0) were euthanized by decapitation. Brains removed from the skull and placed in cold HBSS. The cerebrum was detached from other brain regions under a stereomicroscope by removal of the olfactory bulb and cerebellum, and meninges were peeled off with a tweezer. For cortical neurons, the cortex was dissected out with forceps by removing the hippocampus and the entire midbrain region. For striatal neurons, the hippocampus and the cortex were removed using forceps. The cortex or striatum was minced into approximately 1-2 mm pieces using a sterile razor blade, and digested with 0.05% (w:v) trypsin in HBSS for 10 min at 37°C. Trypsinized tissue was washed three times with HBSS to remove trypsin, and resuspended in DMEM supplemented with 10% FBS (Gibco cat# 2614-079), 100 U/ml penicillin, 100 μg/ml streptomycin (complete neuro DMEM) before passing through a sterile 40 μm cell strainer (Fisherbrand, cat# 22363547) to obtain a cell suspension. Approximately 75,000-100,000 cells were plated on 5 mm diameter coverslips (Word Precision Instruments cat# 502040; coated overnight with 0.1 mg/mL poly-L-lysine hydrobromide [Millipore Sigma cat# P9155], and washed 3x with HBSS) and placed in 96-well plates containing 200 μl of complete neuro DMEM for 4 h, before replacing one half of the volume of medium in each well with Neurobasal medium (GIBCO, cat# 21103049) with B-27 supplement (GIBCO, cat# 17504001) and 1x Glutamax-I (GIBCO, cat# 35050061) (complete neural medium). Approximately 48 h later (day in vitro 3, DIV3), half of the volume of medium in each well was replaced with complete neural media supplemented with 5 μM AraC to achieve a final concentration of 2.5 μM in the well. Beginning DIV5, half of the volume of medium in each well was replaced by fresh complete neural medium every other day.

#### Mouse primary cortical astroglial cell culture

Astrocyte-rich glial cultures were prepared from the cortex of neonatal mice using a protocol similar to that used for cortical neuron cultures except for the following differences. The initial tissue was obtained from neonatal mouse pups (P0-3)^129^, and after straining trypsin-digested tissues, the cell suspension was used to seed 1.5 million cells in each well of a poly-L-lysine coated 6-well plate in complete neuro DMEM. Media was changed the following day, and cells were subsequently split at a 1:2 ratio every 2-3 days by trypsinization followed by centrifugation at 180 x g for 5 minutes before resuspending and reseeding in complete neuro DMEM. Cells were cultured for not more than 5 passages.

#### Mouse primary lung fibroblast cell culture

Lung fibroblasts were isolated from 8-16 week old mice as previously described^130^ with modifications. Mice were euthanized by CO_2_ inhalation followed by cervical dislocation. An incision was made with scissors to open the peritoneal cavity and the skin was removed to expose the cardiothoracic cavity. The sternum was cut open and the ribcage was held open using hemostat forceps. An incision was made in the liver, and DPBS containing calcium and magnesium (Corning, cat# 21-030-CV) was slowly injected into the right ventricle of the heart using a 25-gauge needle to perfuse the lungs. Both lungs and the heart were then dissected out and placed in cold DPBS with calcium and magnesium. The heart and any remaining parts of the trachea or other connective tissue were removed, and the outside of the lungs was rinsed three times in DPBS containing calcium and magnesium. Lungs from 4 mice were minced into ∼1 mm pieces using curved scissors and digested with 1 mg/ml collagenase type 1 (Worthington, cat# LS004194) and 1X dispase (Corning, cat# 354235) in 12 ml of DPBS containing calcium and magnesium for 45 min on a shaker at 37°C. The digested tissue was passed through a 40 μm cell strainer (Fisherbrand, cat# 22363547) to obtain a cell suspension. Twenty ml of DMEM supplemented with 15% FBS, 100 U/ml penicillin, 100 μg/ml streptomycin, and 2 mM L-glutamine (complete fibroblast DMEM) were added to neutralize the digestion cocktail and the tube was spun at 300 x g for 5 min. The pellet was resuspended in 20 ml of complete fibroblast DMEM and cells corresponding to tissue from 4 mice were seeded on two 10 cm tissue culture plates. Cells were washed three times with PBS the following day and fresh complete fibroblast DMEM was added. Two days later, cells were split at a 1:3 ratio by trypsinization followed by centrifugation at 250 x g for 4 minutes before resuspending and reseeding in complete fibroblast media on a total of six 10 cm tissue culture plates. Three days later, all cells were harvested by trypsinization and split in 7 1.2 ml cryogenic vials (Fisherbrand, cat# 1050025) in complete fibroblast media supplemented with 20% FBS and 10% DMSO, and frozen in liquid nitrogen as passage 1 cells for further use. For experiments, these passage 1 cells were quickly thawed in a 30°C water bath, and seeded in a 10 cm culture dish in 15 ml of complete fibroblast media. Media was replaced the day after and cells cultured by splitting as described above for not more than 4 passages.

#### Human primary cardiac fibroblasts

Human cardiac fibroblasts were obtained from ScienCell (cat# 6340) and cultured as previously described^131^. After thawing quickly in a 30°C water bath, ∼350 thousand cells were seeded in each well of a 6-well plate precoated overnight with 0.4% Matrigel (Corning, cat# 356231) in DMEM, and cultured in fibroblast growth medium 3 (Promocell, cat# C-23025) supplemented with 10% growth medium supplement (Promocell, C-39345), 5 μM SB431542 (Selleckchem, cat# S1067), 100 U/ml penicillin, 100 μg/ml streptomycin, and 2 mM L-glutamine. Every two days, cells were split at a 1:2 ratio by incubation in Accutase (Sigma cat# A6964) at 37°C for 3 min to detach cells, followed by centrifugation at 200 x g for 3 min before resuspending and reseeding in fibroblast growth medium 3 supplemented with 10% growth medium supplement, 5 μM SB431542, 100 U/ml penicillin, 100 μg/ml streptomycin, and 2 mM L-glutamine. Cells were typically used within 5 passages, and never cultured for more than 9 passages.

#### Human primary umbilical vein endothelial cells

Human umbilical vein endothelial cells (HUVECs) were acquired from Lonza (cat# CC-2519) and cultured in EBM Basal Medium (Lonza, cat# CC-3121) supplemented with the EGM Singlequots Supplement Pack (Lonza, cat# CC-4133) following the supplier’s recommendations. After thawing quickly in a 30°C water bath, ∼600 thousand cells were seeded on a 10 cm tissue culture dish. Cells were split at a 1:2 ratio every 2 days by trypsinization (Lonza cat# CC-5012), followed by centrifugation at 200 x g for 3 min before resuspending and reseeding in EBM Basal Medium supplemented with the EGM Singlequots Supplement Pack. Cells were cultured for not more than 8 passages.

#### Human primary bronchial smooth muscle cells

Human bronchial smooth muscle cells were obtained from Promocell (cat# C-12561) and cultured in SMC Basal medium 2 (Promocell, cat# C-22262) supplemented with Growth Medium 2 supplement Mix (Promocell, cat# 39267), 100 U/ml penicillin, 100 μg/ml streptomycin, and 2 mM L-glutamine following the supplier’s recommendations. After thawing quickly in a 30°C water bath, ∼200 thousand cells were seeded in each well of a 6 well plate. Cells were split at a 1:2 ratio every 3 days by trypsinization (Promocell cat# C41210), followed by centrifugation at 220 x g for 3 min before resuspending and reseeding in SMC Basal medium 2 supplemented with Growth Medium 2 supplement Mix, 100 U/ml penicillin, 100 μg/ml streptomycin, and 2 mM L-glutamine. Cells were cultured for not more than 5 passages.

### Method Details

#### Plasmids

To generate plasmids encoding putative G protein-binding peptides fused to GST (amino acid sequences presented in **Fig. 1A**), complementary primers encoding the peptides and overhanding ends complementary to EcoRI and HindIII were annealed and inserted between the EcoRI and HindIII site of the previously described pGEX-KG-KB1753 plasmid^132^ to replace the existing KB1753 sequence.

Plasmids encoding KB1691-Nluc (pcDNA3.1-mas-KB1691-Nluc) and KB2123-Nluc (pcDNA3.1-mas-KB2123-Nluc) with a membrane anchoring sequence (mas) were generated by replacing the GRK3ct sequence of pcDNA3.1-mas-GRK3ct-NLuc^27^ by digestion with HindIII/BamHI and subsequent insertion of the KB1691 sequence (SRELAWGISEWLEEWG) or the KB2123 sequence (SSSELADYFGWDGWPG) flanked with a Gly-Gly linker on each end. Plasmids encoding KB1753-Nluc (pcDNA3.1-mas-KB1753-Nluc), GRK2^RH^-Nluc (pcDNA3.1-mas-GRK2^RH^-Nluc), and PRG^RH^-Nluc (pcDNA3.1-mas-PRG^RH^-Nluc) have been described previously^30^. Plasmids encoding human Gαs-67V (pcDNA3.1-Gαs-67V), Gαs-99V (pcDNA3.1-Gαs-99V), and Gαs-154V (pcDNA3.1-Gαs-154V) have previously been described^38^. Plasmids encoding rat Gαi3-YFP (pcDNA3.1-Gαi3-YFP)^113^, mouse Gαq-V (pcDNA3.1-Gαq-V)^96^, and human Gα13-YFP (pcDNA3.1-Gα13-YFP)^30^, all of which are internally tagged with the fluorescent protein at the αb-αc loop of the G protein, have been previously described. The plasmids encoding Gαs/Gαi chimeras were generated using Gibson assembly by swapping either CT-s, αNβ1-s, or both to CT-i and αNβ1-i on human Gαs-99V, and swapping either CT-i, αNβ1-i, or both to CT-s and αNβ1-s on rat Gαi3-YFP. The plasmid encoding FLAG-Ric-8B (pcDNA3.1-FLAG-Ric-8B) was a gift from Dr. Kirill Martemyanov and has been described previously^114^. Plasmids encoding untagged Gβ1 and Gγ2 (pcDNA3.1-Gβ1, pcDNA3.1-Gγ2) have been described previously^26^. The plasmid encoding Glosensor 22F was acquired from Promega (cat# E2301). The plasmid encoding cytosolic KB1691-Nluc (pcDNA3.1-cyto-KB1691-Nluc) was generated by mutation of the glycine in position 2 of pcDNA3.1-mas-KB1691-Nluc to alanine, which removes the myristoylation site of the membrane anchoring sequence. The plasmid encoding for Venus fused to the membrane anchoring sequence of kRas (pVenus-C1-Venus-K-Ras) was a gift from Dr. Nevin Lambert^115^. The pSPAX2 and pMD2.G plasmids for lentiviral packaging were acquired from Addgene (#12260 and #12259).

To generate plasmids encoding ONE-GO biosensors of Gα-GTP species under a CMV promoter, sequences of Gα proteins internally tagged with a YFP at a loop of the all-helical domain along with sequences from an IRES and an Nluc-fused detector module were inserted between the BamHI and MluI sites of vector pLVX-IRES-Hyg by Gibson assembly. The species of the G proteins and amino acid (aa) position of YFP insertion were as follows: Gαs (human, aa99), Gαi1 (rat, aa121), Gαi2 (rat, aa114), Gαi3 (rat, aa114), GαoA (rat, aa119), GαoB (rat, aa119), Gαz (rat, 119), Gαq (mouse, aa124), Gα12 (human, aa140), Gα13 (human, aa131). The IRES used for the ONE-GO biosensors (denoted IRES* in **Fig. S6**) has previously been described^56^, which includes a spacer sequence before the post-IRES start codon and results in diminished expression relative to the coding sequence right downstream of the promoter^56,57^. For Gαs ONE-GO, a T2A-Ric-8B sequence was inserted after KB1691-Nluc (without a stop codon). To generate SV40-Gαi3 ONE-GO and SV40-Gαs ONE-GO constructs, a previously described plasmid (pLVX-mVenus-MYC-DAPLE, based on the pLVX-IRES-Hyg backbone)^133^ was digested with ClaI and SpeI to remove the CMV promoter and insert an SV40 promoter in its place to create pLVX-SV40-mVenus-MYC-DAPLE. The mVenus-MYC-DAPLE cassette between the BamHI and MluI sites was replaced by the same components of the Gαi3 ONE-GO and Gαs ONE-GO constructs described above. All ONE-GO biosensor sequences are provided in Supplemental Text 1. To generate plasmids encoding ONE-GO biosensors of Gα-GTP species under the neuron-specific promoter of human Synapsin (hSyn), the CMV promoter was removed from the plasmids described above by digestion with ClaI and SpeI, followed by insertion of the hSyn promoter followed by a Xho I site in the same sites by Gibson assembly. The sequence of these plasmids (pLVX-hSyn-Gαi1-YFP-IRES*-KB1753-Nluc, pLVX-hSyn-GαoA-YFP-IRES*-KB1753-Nluc, and pLVX-hSyn-Gαq-V-IRES*-GRK2^RH^-Nluc) are provided in Supplemental Text 1.

To generate constructs encoding Gβγ ONE-GO biosensors for different Gα subunits, we first generated a parental plasmid without the Gα component (pLVX-CMV-IRES^WT^-GRK3ct-Nluc-T2A-Gβ1-T2A-Gγ2-V). For this, the vector pLVX-IRES-Hygro was digested with NotI and MluI, and sequences for IRES^WT^, GRK3ct-Nluc, and T2A-Gβ1-T2A-Gγ2-V were inserted by Gibson assembly. In contrast to IRES*, IRES^WT^ does not contain a spacer sequence before the post-IRES start codon and results in comparable expression relative to the coding sequence right downstream of the promoter. To generate plasmids encoding Gαs/Gβγ ONE-GO and Gαi3/Gβγ ONE-GO biosensors, human Gαs or rat Gαi3 were inserted in the NotI site of the parental plasmid described above. In the case of Gαs, a BamHI site was included right before the start codon. Full sequences of parental Gβγ ONE-GO, Gαs/Gβγ ONE-GO and Gαi3/Gβγ ONE-GO biosensor are provided in Supplemental Text 1.

Plasmids encoding GPCRs were obtained from various sources (as indicated Key Resources Table) or subcloned into pcDNA3.1 from the PRESTO-Tango GPCR kit (Addgene Kit #1000000068) as described next. Briefly, GPCR sequences were amplified using primers that preserved the N-terminal FLAG tag present in the Tango library but introduced a stop codon after the end of the sequence of the native GPCR (i.e., it did not include the V2-tail, TEV site and tTA transcription factor present in the Tango constructs). The amplicons were inserted between the NotI and Apa I sites of pcDNA3.1 by Gibson assembly. The FLAG tag was not included for the following mGluR-encoding plasmids generated from the PRESTO-Tango library: pcDNA3.1(+)-GRM1, pcDNA3.1(+)-GRM2, pcDNA3.1(+)-GRM5, pcDNA3.1(+)-GRM7, and pcDNA3.1(+)-GRM8.

All point mutations were generated using QuikChange II following the manufacturer’s instructions (Agilent, 200523).

#### Pulldown assays

Putative G protein-binding peptides fused to GST were expressed in BL21(DE3) *E. coli* transformed with the corresponding plasmids by overnight induction at 23°C with 1 mM isopropyl β-D-1-thio-galactopyranoside (IPTG). IPTG was added when the OD600 reached ∼0.8. Bacteria pelleted from 50 ml of culture were resuspended at 4°C in 10.5 ml of lysis buffer (50 mM NaH_2_PO_4_, pH 7.4, 300 mM NaCl, 10 mM imidazole supplemented with a protease inhibitor mixture of 1 μM leupeptin, 2.5 μM pepstatin, 0.2 μM aprotinin, and 1 mM phenylmethylsulfonyl fluoride). The cell suspension was supplemented with 1% (v/v) Triton X-100 and sonicated (3 pulses of 20 s separated by 30 s intervals for cooling). The resulting lysate was cleared by centrifugation at 4,300 × g for 40 min at 4°C. The supernatant for each construct was collected and 2 ml aliquots were frozen until use.

For pulldown assays, an aliquot of lysate containing ∼25-50 μg of each GST-fused peptide was incubated with ∼20 μl of glutathione-agarose beads (Thermo, 16100) at 4°C for 90 min. Beads were washed twice with lysis buffer and resuspended in 250 μl of binding buffer (50 mM Tris-HCl, 100 mM NaCl, 0.4% NP-40, 10 mM MgCl_2_, 5 mM EDTA, 1 mM DTT) supplemented with 30 μM GDP, 30 μM AlCl_3_, and 10 mM NaF. His-tagged Gαs or Gαo, purified as described previously^134,135^, was diluted in binding buffer supplemented with 30 μM GDP, 30 μM AlCl_3_, and 10 mM NaF to obtain 2 μg of protein in 50 μl, and incubated at 30°C for 60 min. GDP-AlF ^-^-loaded G proteins were centrifuged at 14,000 x g for 3 min before addition to tubes containing the GST-fused peptides immobilized on glutathione-agarose beads. Tubes were incubated for 4 h at 4°C with constant rotation. Beads were washed three times with 1 ml of wash buffer (4.3 mM Na_2_HPO_4_, 1.4 mM KH_2_PO4, pH 7.4, 137 mM NaCl, 2.7 mM KCl, 0.1% (v/v) Tween 20, 10 mM MgCl_2_, 5 mM EDTA, 1 mM DTT) supplemented with 30 μM GDP, 30 μM AlCl_3_, and 10 mM NaF, and resin-bound proteins were eluted with Laemmli sample buffer by incubation at 37°C for 10 min. Proteins were separated by SDS-PAGE and immunoblotted with antibodies as indicated under “*Protein Electrophoresis and Immunoblotting*.”

#### Bioluminescence Resonance Energy Transfer (BRET) measurements in HEK293T cells

HEK293T cells were seeded on 6-well plates (∼400,000 cells/well) coated with 0.1% gelatin (Sigma-Aldrich cat# G1393), and transfected ∼24 h later using the calcium phosphate method. For experiments with bi-molecular biosensors consisting of YFP-tagged Gα proteins and Nluc-fused detector modules (**Fig. 1, 2, 3**; **Fig. S1, S3, S4, S5, S6**), cells were transfected in the combinations indicated in the figures with the following plasmids (amount per well indicated in parenthesis): Gαs-67V (1 μg), Gαs-99V (1 μg), Gαs-154V (1 μg), Gαi3-YFP (1 μg), Gαq-V (1 μg) or Gα13-YFP (1 μg), KB1691-Nluc (0.05 μg), KB2123-Nluc (0.05 μg), KB1753-Nluc (0.05 μg), GRK2^RH^-Nluc (0.05 μg), PRG^RH^-Nluc (0.05 μg), Ric-8B (0.1 μg), Gβ1 (0.2 μg), Gγ2 (0.2 μg), and different GPCRs (0.2 μg). For the bystander Gαs biosensor system (**Fig. S2**), cells were transfected with the following plasmids (amount per well indicated in parenthesis): Gαs (0.5 μg), cytosolic KB1691-Nluc (0.01 μg), Venus-kRas (0.25 μg), and β2AR (0.2 μg). Unless otherwise indicated in the figures, the amount of plasmid transfected per well for Gα-GTP ONE-GO biosensors and Gβγ ONE-GO biosensors was 0.05 μg and 1μg, respectively, and the amount of GPCR-encoding plasmids co-transfected was 200 ng, except for plasmids encoding A1R, P2Y6 and P2Y12 in **Fig. 7**, which was 5 ng to reduce basal antivity.

Six hours after transfection, media was replaced by fresh complete medium or medium with reduced serum (0.1% FCS) in the case of experiments with glutamate receptors (**Fig. 4B; Fig. S7**). Cells were harvested 18-24 h later for luminescence measurements by washing with PBS once and gentle scraping in 1 ml of PBS. Cells were centrifuged for 5 min at 550 x g, and resuspended in BRET buffer (140 mM NaCl, 5 mM KCl, 1 mM MgCl_2_, 1 mM CaCl_2_, 0.37 mM NaH_2_PO_4_, 20 mM HEPES pH 7.4, 0.1% glucose) at a concentration of approximately 1 million cells/ml. Twenty-five to fifty thousand cells were added to a white opaque 96-well plate (Opti-Plate, PerkinElmer cat# 6005290) and mixed with the nanoluciferase substrate Nano-Glo (Promega cat# N1120, final dilution 1:200) for 2 min before measuring luminescence in a POLARstar OMEGA plate reader (BMG Labtech) at 28°C. Luminescence was measured at 460 ± 40 nm and 535 ± 15 nm, and BRET signal was calculated as the ratio between the emission intensity at 535 nm divided by the emission intensity at 460 nm.

For kinetic BRET measurements, luminescence signals were measured every 0.24 s (with a signal integration time of 0.24 s) for the duration of the experiment. Reagents were added to the wells during live measurements using injectors. BRET data are presented as the difference from baseline BRET signal [ΔBRET (baseline)] by subtracting the average of signal pre-stimulation from all data points. A similar procedure with minor modifications was followed for time-resolved multi-well experiments (**Fig. S8C**). Briefly, cells were added to 48 wells of an assay plate and incubated with CTZ400a as the luciferase substrate (GoldBio cat# C-320-1; final concentration of 10 μM) before measuring luminescence in each well every 15 seconds. The Z’ values for the time-resolved multi-well experiments were calculated as described previously ^136^ by using the formula Z’ = 1−[3*(δ_positive_ + δ_negative_)/│µ_positive_ - µ_negative_│], where δ is the standard deviation, µ is the mean, and positive / negative indicates presence of absence of agonist, respectively.

For endpoint BRET measurements to determine dose dependence responses, CTZ400a (final concentration of 10 μM) was used as luciferase substrate, and the BRET signal (535 nm luminescence / 460 nm luminescence) was measured every minute for 5 minutes with a signal integration time of 0.32 s for each measurement. Calculations were done with data corresponding to BRET signals at different time points depending on the kinetics of the response for different receptors and G proteins. BRET data are presented as the difference from BRET signal relative to a condition without agonist [ΔBRET (no agonist)]. Where indicated, the EC_50_ and pEC_50_ values were determined by using a 3-parameter sigmoidal curve-fit in Prism (GraphPad). To compare EC_50_ values experimentally determined with ONE-GO biosensors for different GPCRs and existing curated pharmacological parameters (**Fig. 4B**), we extracted pK_d_, pK_i_ or pE/IC_50_ values for from the IUPHAR database^61^, with the following prioritization based on availability : pK_d_ > pK_i_ > pE/IC_50_. The data was extracted for GPCRs of the same species as the ones used in the experiments with ONE-GO sensors, and when a range of values were given the midpoint was used. For some experiments, the data was further processed in the following ways. For **Fig. 2C, D** the ΔBRET values for each G protein chimera (shown in **Fig. S4**) were normalized by using the maximal ΔBRET value obtained with the cognate wild-type G protein for each GPCR as 1. For **Fig. 5A**, the ΔBRET values were normalized by using the ΔBRET value obtained upon stimulation with an EC_80_ concentration of the cognate agonist for each GPCR as 1. Data processed this way was used to generate the Principal Component Analysis (PCA) presented in this panel using the PCA tool with the centered method and with all principal components selected in Prism (GraphPad). For **Fig. S8A**, the ΔBRET values were normalized by using the ΔBRET value obtained upon stimulation with an EC_100_ concentration of the cognate agonist for each wild-type GPCRs as 1. For **Fig. 5C**, the ΔBRET values were first corrected by subtraction of BRET values obtained with cells treated with the same ligands but without GPCR transfection (shown in **Fig. S8B**), followed by normalization using the corrected ΔBRET value corresponding to the maximal response observed for a given G protein across all three GPCR as 1. For curves in **Fig. S8B** with detectable responses (> 0.001 ΔBRET) that could not be reliably fit to accurately determine EC_50_ values, an arbitrary EC_50_ value of 30 mM was given.

At the end of some BRET experiments, a separate aliquot of the same pool of cells used for the luminescence measurements was centrifuged for 1 min at 14,000 x g and pellets stored at −20°C for subsequent immunoblot analysis (see “*Protein electrophoresis and Immunoblotting*” section below).

#### Luminescence-based cAMP measurements in cells

Approximately 300,000 HEK293T cells were seeded on each well of 6-well plates coated with 0.1% (w/v) gelatin, and transfected with plasmids ∼24 hr later using the calcium phosphate method. For experiments testing the effect of the multi-plasmid Gαs sensor system on cAMP signaling (**Fig. 1D**), cells were transfected with the following plasmids (amount per well indicated in parenthesis): Gαs-99V (1 μg), FLAG-Ric-8B (0.1 μg), Gβ1 (0.2 μg), Gγ2 (0.2 μg), β2AR (0.2 μg), with or without KB1691-Nluc (0.05 μg) and supplemented with pcDNA3.1 to equalize total amount of DNA per well. For experiments testing the effect of Gα ONE-GO biosensors on cAMP signaling (**Fig. 3D**), cells were transfected with the following plasmids (amount per well indicated in parenthesis): Gαs or Gαi3 ONE-GO biosensor (0.05 μg), α2_A_-AR (0.2 μg; only in combination with Gαi3 ONE-GO) and supplemented with pcDNA3.1 to equalize total amount of DNA per well. The amount of Glosensor 22F plasmid transfected per well for all experiments was 0.8 μg. Cell medium was changed 6 h after transfections.

Approximately 16-24 h after transfection, cells were washed and gently scraped in room temperature PBS, and centrifuged (5 min at 550 × g). For kinetic measurements, cells were resuspended in 750 μl Tyrode’s buffer (140 mM NaCl, 5 mM KCl, 1 mM MgCl_2_, 1 mM CaCl_2_, 0.37 mM NaH_2_PO_4_, 24 mM NaHCO_3_, 10 mM HEPES and 0.1% glucose, pH 7.4). Two-hundred μl of cells were mixed with 200 μl of 5 mM D-luciferin K+ salt (GoldBio, LUCK-100) diluted in Tyrode’s buffer and incubated at 28 °C for 15 minutes. Ninety μl of cells pre-incubated with D-luciferin were added to a white opaque 96-well plate (Opti-Plate, PerkinElmer Life Sciences, 6005290) before measuring luminescence without filters at 28 °C every 10 s in a BMG Labtech POLARStar Omega plate reader. Agonists were added as indicated in the figures during the recordings using built-in injectors. Kinetic traces are represented as raw luminescence signal.

For dose response experiments, cells were washed and scraped as above, but resuspended in 300 μl Tyrode’s buffer. Two-hundred and forty μl of cells were mixed with 240 μl of 5 mM D-luciferin K+ salt diluted in Tyrode’s buffer and incubated at 28 °C for 15 minutes. For experiments testing the effect of Gαs ONE-GO expression, 20 μl of different doses of isoproterenol diluted in Tyrode’s buffer at 5X the final concentration desired in the assay were added to wells of a white opaque 96-well plate and further diluted with. 57.6 μl of Tyrode’s buffer. Reactions were initiated by addition of 22.4 μl of the suspension of cells pre-incubated with D-luciferin and luminescence measurements were immediately started. Luminescence signal was measured without filters at 28 °C every 30 s in a BMG Labtech POLARStar Omega plate reader. For each dose of isoproterenol, the response values were calculated by averaging the 3 time points around the peak of the kinetic trace (270, 300, and 330 s after start of measurement) and normalizing them by using the response at maximal dose of isoproterenol as 100. For experiments testing the effect of Gαi3 ONE-GO expression, 20 μl of different amounts of brimonidine diluted in Tyrode’s buffer at 4X the final concentration desired in the assay were added to wells of a white opaque 96-well plate, and further diluted by addition of 37.6 μl of Tyrode’s buffer. Reactions were initiated at room temperature by addition of 22.4 μl of the cell suspension pre-incubated with D-luciferin, and 2 minutes later 20 μl of 500 nM isoproterenol (to achieve a final concentration of 100 nM in the assay) diluted in Tyrode’s buffer were added, followed immediately by luminescence measurements as was done above for Gαs ONE-GO. For each dose of brimonidine, response values were calculated by averaging the 3 time points around the peak of the kinetic trace (270, 300, and 330 s after start of measurement) and normalizing them by using the response in the absence of brimonidine as 100.

At the end of some experiments, a separate aliquot of the same pool of cells used for the measurements was centrifuged for 1 min at 14,000 x g and pellets stored at −20°C for subsequent immunoblot analysis (see “*Protein electrophoresis and Immunoblotting*” section below).

#### BRET measurements in HeLa cells

HeLa cells stably expressing Gαi3 ONEGO or Gαi3/Gβγ ONEGO were generated by lentiviral transduction followed by Fluorescence-Activated Cell Sorting (FACS). Lenti-X 293T (Cat# 632180, Takara Bio) cells were seeded on 6-well plates (∼400,000 cells/well) coated with 0.1% gelatin and transfected the next day using the polyethylenimine (PEI) method^137^ at a 2:1 PEI:DNA ratio with the following plasmids (amount of DNA per well in parenthesis): pLVX-CMV-Gαi3-YFP-IRES*-KB1753-Nluc or pLVX-CMV-Gαi3-IRES^WT^-GRK3ct-Nluc-T2A-Gβ1-T2A-Gγ2-V (1.8 µg), psPAX2 (1.2 µg), and pMD2.G (0.75 µg). The DNA was added to 200 μl of DMEM and mixed with 7.5 μl of PEI reagent (Polysciences Inc cat# 23966). Tubes were incubated at room temperature for 15 min before adding to cells. Six hours after transfection, media was replaced with DMEM supplemented with 10% FCS, 100 U/ml penicillin, 100 μg/ml streptomycin, and 2 mM L-glutamine (complete medium). Lentivirus-containing media was collected 24 hr and 48 hr after transfection, centrifuged at 1500 x g for 5 min after pooling, and filtered through a 0.45-μm surfactant-free cellulose acetate (SFCA) membrane filter (Corning cat# 431220).

HeLa cells were seeded on 6-well plates (∼200,000 cells per well) precoated with 0.1 mg/mL poly-L-lysine hydrobromide (overnight incubation followed by 3 washes with HBSS) and transduced the next day by a 48 hr incubation in 2 ml of a 1:1 mix of lentivirus-containing supernatants described above mixed with fresh complete media, supplemented with 6 µg/ml of polybrene (Tocris Bioscience cat# 7711/10). Cells were first transferred from the 6-well plate to a 10-cm plate after virus transduction for expansion. Once the 10-cm plate was confluent, cells were further expanded into 15-cm dishes to have enough material for sorting as described next. For FACS, HeLa cells were trypsinized, resuspended in complete medium, and 7.5 million cells were transferred to a 15 ml conical tube. Cells were washed 3 x with 10 ml cold PBS by cycles of centrifugation (300 x g for 3 minutes), aspiration, and resuspension. Cells were resuspended in 1.5 ml cold PBS and stored on ice until sorting. A subset of the trypsinzed HeLa cells were resuspended in complete DMEM containing DAPI (1 μg/ml), washed as described above, and used for the gating process. Cell sorting was performed on a Moflo Astrios EQ (Beckman Coulter), and the 488_ex_/513_em_ nm fluorescence channel (Voltage: 354 nV) was used for positive selection. Cells with fluorescent intensity above 300 were collected as “*High sensor expression*”, and cells with fluorescent intensity from 80 to 300 were collected as “*Low sensor expression*”. Cells were collected and seeded in a 6-well plate with complete DMEM for expansion.

For BRET experiments, HeLa cells stably expressing Gαi3 ONEGO or Gαi3/Gβγ ONEGO were seeded on 5 mm glass coverslips precoated with 0.1 mg/mL poly-L-lysine hydrobromide (overnight incubation followed by 3 washes with HBSS) and placed in a 96-well plate (∼35,000 cells per well). Recordings were done 24 h after seeding as described next. Coverslips were washed with 200 μl BRET buffer and transferred to a well of a white opaque 96-well plate containing BRET buffer and Nano-Glo (final dilution 1:200) with tweezers, followed by incubation in the dark for 2 min at room temperature before measuring luminescence in a POLARstar OMEGA plate reader (BMG Labtech). Measurements and data processing were done as described in “*BRET measurements in HEK293T cells*” except that measurements with Gαi3/Gβγ ONEGO were made every 0.48 s (with a signal integration time of 0.48 s).

To collect samples for immunoblotting, HeLa cells were seeded on a 6-well plate as described above, scraped in PBS and spun at 550 x g for 5 min. Pellets were processed for immunoblotting as described below in “*Protein electrophoresis and Immunoblotting*”.

#### Production of concentrated lentiviral particles

Lentiviruses used for transduction of SH-SY5Y cells and primary cells were concentrated after large scale packaging as described previously^137,138^. Lenti-X 293T cells were plated on 150 mm diameter dishes (∼2.5 million cells / dish) and cultured at 37°C, 5% CO_2_ in DMEM supplemented with 10% FCS, 100 U/ml penicillin, 100 μg/ml streptomycin, and 2 mM L-glutamine. After 16-24 h, cells were transfected using the polyethylenimine (PEI) method^137^ at a 2:1 PEI:DNA ratio with the following plasmids (amount of DNA per dish in parenthesis): psPAX2 (18 μg), pMD2.G (11.25 μg) and Gα ONE-GO biosensor (27 μg). Approximately 16 h after transfection, media was replaced. Lentivirus containing media was collected 24 and 48 h after the initial media change (∼70 mL per dish and 4 dishes for each construct). Media was centrifuged for 5 min at 900 x g and filtered through a 0.45 μm sterile PES filter (Fisherbrand cat# FB12566505). Filtered media was centrifuged for ∼18 h at 17,200 x g at 4°C (Sorvall RC6+, ThermoScientific F12-6x500 LEX rotor) to sediment lentiviral particles. Pellets were washed and gently resuspended in 1 mL of PBS and centrifuged at 50,000 x g for 1 h at 4°C (Beckman Optima MAX-E, TLA-55 rotor). Pellets were resuspended in 500 μl of PBS to obtain concentrated lentiviral stocks that were stored at −80°C in aliquots. Each aliquot was thawed only once and used for less than a week stored at 4°C for subsequent experiments.

#### BRET measurements in SH-SY5Y cells

Naïve SH-SY5Y cells and SH-SY5Y cells stably expressing Gαi*-BERKY3^30^ were seeded on 35 mm plates (∼700,000 cells/plate) coated with 0.1% gelatin. The following day, naïve cells were transduced by replacing the media with fresh medium supplemented with concentrated lentiviral particles of the Gαi3 ONE-GO biosensor under the CMV promoter (1:400 final dilution). Approximately 18 h later, lentivirus-containing media was replaced with fresh medium. Both SH-SY5Y cells stably expressing Gαi*-BERKY3 and cells expressing the Gαi3 ONE-GO sensor were harvested for luminescence measurements approximately 48 h after plating.

Briefly, cells were washed with PBS once, scraped, and spun at 550 x g for 5 min. Cells were resuspended in BRET buffer at a concentration of approximately 1 million cells/ml. For luminescence measurements, 40,000-100,000 cells were added to a white opaque 96-well plate and kinetic BRET measurements were performed as described in “*Bioluminescence Resonance Energy Transfer (BRET) measurements in HEK293T cells*”. At the end of some BRET experiments, a separate aliquot of the same pool of cells used for the luminescence measurements was centrifuged for 1 min at 14,000 x g and pellets stored at −20°C for subsequent immunoblot analysis (see “*Protein electrophoresis and Immunoblotting*” section below).

#### BRET measurements in primary cell cultures

ONE-GO biosensors were expressed in primary cells by lentiviral transduction according to the specific procedures described below using concentrated stocks described in “*Production of concentrated lentiviral particles*”. All primary cells were transduced with lentiviral constructs expressing the ONE-GO biosensors under the CMV promoter, except for neurons, where ONE-GO biosensors were expressed under the human synapsin I (hSyn) promoter. Cell seeding, viral dilutions, and other parameters are detailed next for each one of the cell types investigated. <u>Mouse lung fibroblasts</u> were seeded on 5 mm glass coverslips precoated with 0.1% gelatin (5 min incubation at room temperature followed by aspiration) and placed in a 96-well plate (15,000 cells per well). Approximately 18 h after seeding, cells were transduced by replacing the media with 100 μl of fresh media supplemented with 6 μg/ml polybrene and the following lentiviruses (final dilution in parenthesis): Gαq ONE-GO (1:500), Gαi3 ONE-GO (1:500), Gα13 ONE-GO (1:500). Plates were spun at 600 x g for 30 min and returned to the incubator. Media was replaced ∼24 h later. BRET recordings were done 48 or 72 h after the addition of virus as described below. <u>Mouse astroglial cells</u> were seeded on 5 mm glass coverslips precoated with 0.1 mg/mL poly-L-lysine hydrobromide (overnight incubation followed by 3 washes with HBSS) and placed in a 96-well plate (40,000 cells per well). Approximately 18 h after seeding, cells were transduced by replacing the media with 100 μl of fresh media supplemented with 6 μg/ml polybrene and the following lentiviruses (final dilution in parenthesis): Gαq ONE-GO (1:250), Gαi3 ONE-GO (1:500), Gα13 ONE-GO (1:100-1:250). Plates were spun at 600 x g for 30 min and returned to the incubator. Media was replaced ∼24 h later. BRET recordings were done ∼48 h post-transduction as described below. <u>Mouse cortical and striatal neurons</u> were seeded as explained above in “*Mouse primary cortical & striatal neuron culture*”. On DIV7, cells were transduced by replacing half of the media with fresh media containing the following lentiviruses (final dilution in parenthesis): Gαq ONE-GO (1:200), Gαi1 ONE-GO (1:200), GαoA ONE-GO (1:200). Half of the media was replaced with fresh medium ∼2 h later and BRET recordings were done on DIV12-19 as described below. <u>Human bronchial SMCs</u> were seeded on 5 mm glass coverslips precoated with 0.1% gelatin and placed in a 96-well plate (15,000 cells per well). Approximately 18 h after seeding, cells were transduced by replacing the media with 100 μl of fresh media supplemented with 6 μg/ml polybrene and the following lentiviruses (final dilution in parenthesis): Gαq ONE-GO (1:1,500), Gαs ONE-GO (1:1,500), Gα13 ONE-GO (1:1,500). Plates were then spun at 600 x g for 30 min and returned to the incubator. Media was replaced ∼24 h later, followed by BRET recordings that same day as described below. <u>HUVECs</u> were seeded on 5 mm glass coverslips precoated with 0.1% gelatin and placed in a 96-well plate (15,000 cells per well). Approximately 18 h after seeding, cells were transduced by replacing the media with 100 μl of fresh media supplemented with 6 μg/ml polybrene and the following lentiviruses (final dilution in parenthesis): Gαq ONE-GO (1:2,000), Gαs ONE-GO (1:500), Gαi3 ONE-GO (1:2,000), Gα13 ONE-GO (1:500). Plates were then spun at 600 x g for 30 min and returned to the incubator. Media was replaced ∼24 h later, followed by BRET recordings the same day as described below. <u>Human cardiac fibroblasts</u> were seeded on 5 mm glass coverslips precoated overnight with 0.4% Matrigel in DMEM and placed in a 96-well plate (13,000 cells per well). Approximately 18 h after seeding, cells were transduced the following ways depending on experimental purposes. For **Fig. 6** and **Fig. S10**, cells were washed twice with HBSS to remove SB431542.Then, cells were incubated in 100 μl of fresh media supplemented with 6 μg/ml polybrene and the following lentiviruses (final dilution in parenthesis): Gαq ONE-GO (1:4,000-1:5,000), Gαs ONE-GO (1:1,000), Gαi3 ONE-GO (1:2,000 for adenosine stimulated cells, 1:4,000-1:5,000 for TRAP-6 stimulated cells), Gα13 ONE-GO (1:4,000-1:5,000). Plates were spun at 600 x g for 30 min, returned to the incubator and media was replaced ∼24 h later, followed by BRET recordings 2 h later when testing the effect of adenosine or 24 h later when testing the effect of TRAP-6. For **Fig. 7A**, cells were kept in media supplemented with 5 μM SB431542 throughout. Approximately 18 h after seeding, cells were incubated in 100 μl of fresh media supplemented with 6 μg/ml polybrene and the following lentiviruses (final dilution in parenthesis): Gαq ONE-GO (1:2,000), Gαi3 ONE-GO (1:2,000), Gα13 ONE-GO (1:2,000). Plates were spun at 600 x g for 30 min, returned to the incubator and media was replaced ∼24 h later, followed by BRET recordings 24 h later. For **Fig. 7C**, cells were washed twice with HBSS to remove SB431542 approximately 18 h after seeding. Then, cells were incubated in 100 μl of fresh media supplemented with 6 μg/ml polybrene and the following lentiviruses in the presence or absence of TGFβ (final dilutions without/with 1 nM TGFβ respectively in parenthesis): Gαq ONE-GO (1:2,000/1:5,000), Gαi3 ONE-GO (1:2,000/1:5,000), Gα13 ONE-GO (1:2,000/1:5,000). Plates were spun at 600 x g for 30 min, returned to the incubator and media was replaced ∼48 h later, followed by BRET recordings 2 h later.

Kinetic BRET recordings were done the same way for all cell types. Briefly, coverslips were washed with 200 μl BRET buffer and transferred to a well of a white opaque 96-well plate containing BRET buffer and Nano-Glo (final dilution 1:200) with tweezers, followed by incubation in the dark at room temperature for 2 min before measuring luminescence in a POLARstar OMEGA plate reader (BMG Labtech). Measurements and data processing were done as described in “*BRET measurements in HEK293T cells*” except that measurements were taken every 0.96 s (with a signal integration time of 0.96 s). Where indicated in the figures or figure legends, cells were pretreated with pertussis toxin (overnight), YM-254890 (2 min), or a GPCR antagonist (2 min). To determine “Max ΔBRET” (**Fig. 7A, 7C**), 9 data points centered around the maximum response were averaged. For some representations in **Fig. 7**, Max ΔBRET values were normalized by using the Max ΔBRET value obtained with Gαq ONE-GO for each cell type as 1. The “*Basal BRET*” and “*Luminescence*” values reported in **Fig. 7C** were obtained by averaging the first 10 data points of 535/460 ratios or direct counts in the 460 nm channel, respectively.

For endpoint BRET measurements to determine dose-dependent responses with cardiac fibroblasts (**Fig. 6**), coverslips were washed and transferred to an assay plate as explained above. A 10X agonist solution was simultaneously added to all wells using a multi-channel pipette, and BRET signal (535 nm luminescence / 460 nm luminescence) was measured every minute for 5 minutes with a signal integration time of 0.96 s for each measurement. Calculations were done with data corresponding to BRET signals at different time points depending on the kinetics of the response for different receptors and G proteins. BRET data are presented as the difference from BRET signal relative to a condition without agonist [ΔBRET (no agonist)]. The Gαi3 ONE-GO dose response curve was generated by doing kinetic BRET measurements as explained above and taking the difference in ΔBRET at max amplitude for each dose compared to BRET buffer only injections.

To collect samples of human cardiac fibroblasts for immunoblotting, cells were seeded on a glass bottom 12-well plate precoated overnight with 0.4% Matrigel in DMEM (∼140,000 cells per well). Transductions and TGFβ treatments were done exactly as described above for the BRET measurements. Cells were washed with PBS and 90 μl of ice-cold lysis buffer was added to each well before proceeding with the rest of the steps described in “*Protein electrophoresis and Immunoblotting*” section below.

#### RT-qPCR of cardiac fibroblast mRNA

Human cardiac fibroblasts were seeded on a glass-bottom 12-well plate precoated overnight with 0.4% Matrigel in DMEM (∼140,000 cells per well). Approximately 18 h after seeding, cells were washed 2 x with HBSS, and then incubated in 1 ml of fresh media without SB431542 supplemented (or not) with 1 nM TGFβ. Approximately 48 h later, RNA was extracted using the PureLink RNA Mini kit (ThermoFisher cat# 12183018A) and stored at -80°C. Two-hundred and fifty ng of RNA were used for reverse transcription reactions using the LunaScript RT SuperMix (New England BioLabs cat# E3010). For the qPCR, 10 μl reactions were prepared in technical triplicate for each condition by mixing 1 μl of reverse-transcribed DNA, 4 μl of primer mix (to achieve a final concentration of 10 μM), and 5 μl of 2X SYBR Green PCR Master Mix (Applied biosystems cat# 4385612). For each gene target, the primer mixes consisted of the following forward (F) and reverse (R) primers: h18S (F: CGGCGACGACCCATTCGAAC; R: GAATCGAACCCTGATTCCCCGTC); hACTA2 (F: AAAAGACAGCTACGTGGGTGA; R: GCCATGTTCTATCGGGTACTTC); hCOL1A (F: ACGAAGACATCCCACCAATCACCT; R: AGATCACGTCATCGCACAACACCT); hFN1 (F: GGCAACGTGTTACGATGAT; R: CGGGAATCTTCTCTGTCAG). The qPCR was run on a ViiA 7 Real-Time PCR machine (Applied biosystems) using the following conditions: Initial denaturation was done at 95°C for 20 s, then 40 cycles were repeated going from 1 s at 95°C to 20 s at 60°C.

Relative mRNA expression was calculated using the 2^(-ΔΔCt)^ method as described^139^ using h18S as the housekeeping gene. The fold increase mRNA upon TGFβ treatment was calculated by dividing each value from the previous step by the average mRNA expression of the control condition (without TGFβ) for each target gene.

#### Protein electrophoresis and immunoblotting

Pellets of HEK293T, SH-SY5Y cells, or HeLa cells were resuspended on ice with lysis buffer (20 mM HEPES, 5 mM Mg(CH_3_COO)_2_, 125 mM K(CH_3_COO), 0.4% (v:v) Triton X-100, 1 mM DTT, 10 mM β-glycerophosphate, 0.5 mM Na_3_VO_4_, supplemented with a protease inhibitor cocktail [SigmaFAST cat# S8830], pH 7.4) and thoroughly vortexed. Cardiac fibroblasts were lysed directly in the culture wells as described in “*BRET measurements in primary cell cultures*” with the same buffer and transferred to 1.5 ml tubes before thorough vortexing. Lysates were cleared by centrifugation (10 min at 14,000 x g, 4°C) and quantified by Bradford (Bio-Rad, cat#5000205). All samples were then boiled for 5 min in Laemmli sample buffer, except those used to detect PAR1 (**Fig. 7C**), which were incubated at 37°C for 10 min.

Proteins were separated by SDS-PAGE, followed by transfer to PVDF membranes (EMD Millipore cat# IPFL00010) for 2 h. Membranes were stained with Ponceau S solution (0.1% w/v Ponceau S in 5% v/v acetic acid) and imaged on a flatbed scanner prior to immunoblotting. PVDF membranes were blocked with TBS supplemented with 5% (w;v) non-fat dry milk for 1 hour, and then incubated sequentially with primary and secondary antibodies diluted in TBST supplemented with 2.5% (w:v) non-fat dry milk. Primary antibody species, vendors, and dilutions were as follows (in parenthesis): His (mouse, Sigma cat# H1029; 1:2,500); β-actin (rabbit, LI-COR cat# 926-42210; 1:2,500); GFP (mouse, Clontech cat# 632380; 1:2,000); FLAG (mouse, Millipore Sigma cat# F1804; 1:1000); Gαs (rabbit, SCBT cat# sc-383; 1:500); tubulin (mouse, Sigma cat# T6074; 1:2,500); Gβ (mouse, SCBT cat# sc-166123; 1:250); PAR1 (mouse, Millipore Sigma cat# MABF244; 1:500); Gαi3 (rabbit, Aviva cat# OAAB19207; 1:250), except in **Fig. 3C, D**, which has rabbit, Millipore Sigma cat# 06-270; 1:500; Nluc in **Fig. 1D**, **Fig. S1A-B**, **Fig. S6A**, **Fig. S9** (rabbit, Promega^41^; 1:1,000); Nluc in **Fig. 2A, 3B, 4A, 7C** (mouse, Promega cat# N700A; 1:500-1:1000); Nluc in **Fig. 3C-D** (mouse, R&D systems cat# MAB10026; 1:250). The following secondary antibodies were used at a 1:10,000 dilution (species and vendor indicated in parenthesis): anti-rabbit Alexa Fluor 680 (goat, Invitrogen cat# A21077); anti-mouse Alexa Fluor 680 (goat, Invitrogen cat# A21058); anti-mouse IRDye 800 (goat, LI-COR cat# 926-32210); anti-rabbit DyLight 800 (goat, Thermo cat# 35571). Infrared imaging of immunoblots was performed according to manufacturer’s recommendations using an Odyssey CLx infrared imaging system (LI-COR Biosciences). Images were processed using Image Studio software (LI-COR), and assembled for presentation using Photoshop and Illustrator software (Adobe).

#### Statistical analysis and structure display

Unless otherwise indicated, all experiments were performed at least three times, and presented as averages ± the standard error of the mean (SEM). For the sake of clarity, in the presentation of kinetic BRET measurements the SEM is only presented in the positive direction and displayed as bars of a lighter color tone than that of its corresponding data points. Protein structure images displayed in **Fig. 2A**, **2B** were prepared in ICM (Molsoft LLC., San Diego, CA). Chemical structures (**Fig. 5A, C**) were generated using ChemDraw (PerkinElmer informatics, Waltham, MA).

---

**Figure S1.**
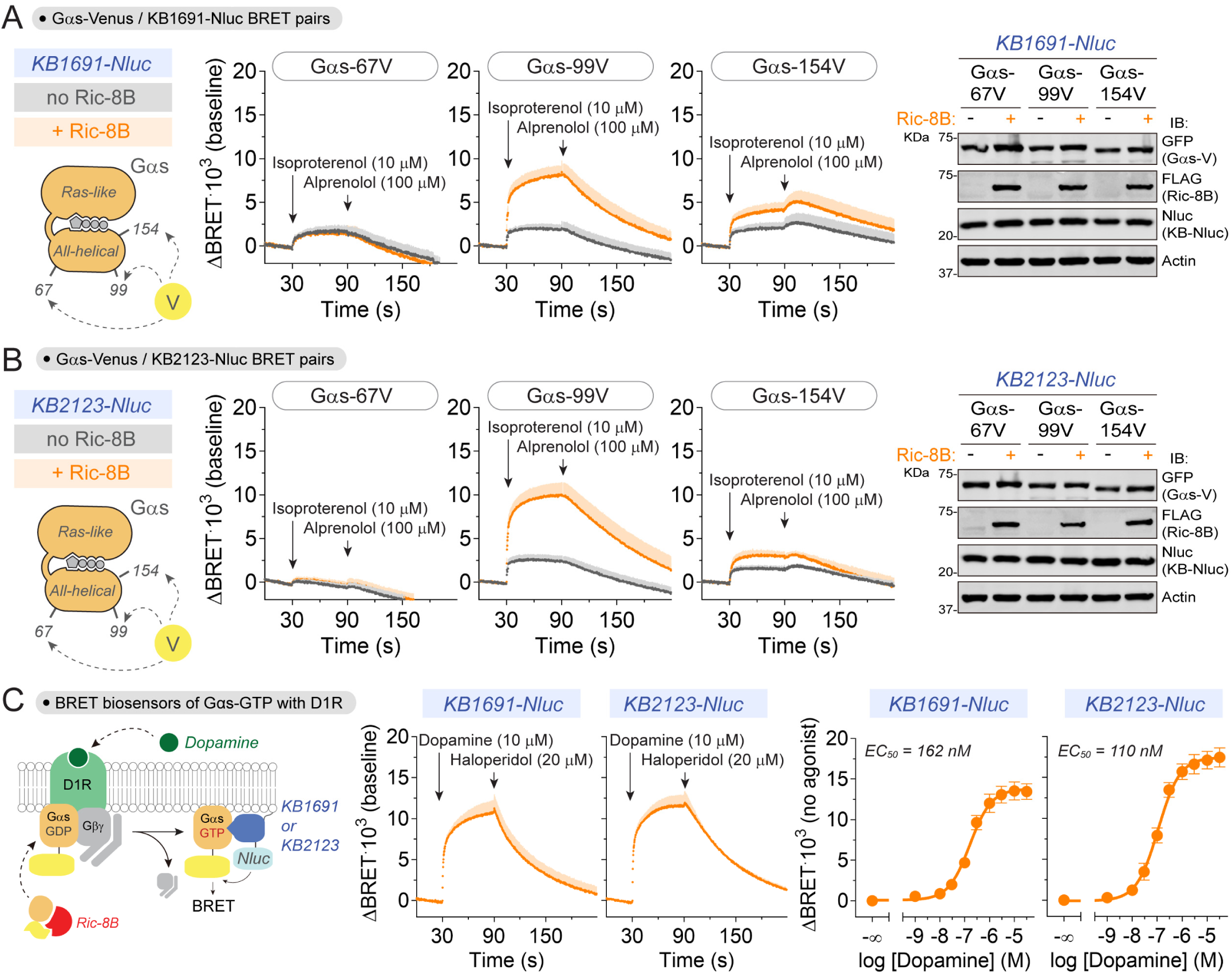
Live-cell Gαs activity biosensor design. **(A, B)** Identification of optimal acceptor design for Gαs-GTP biosensors based on KB1691-Nluc and KB2123-Nluc. BRET responses were measured in HEK293T cells expressing KB1691-Nluc or KB2123-Nluc along with Gβ1, Gγ2, Gαs (short isoform) tagged with Venus at three different internal positions (67V, 99V, or 154V), and with (orange) or without (black) co-expression of Ric-8B. Results are the mean ± S.E.M. of n=3-4. A representative immunoblot showing the expression of the different components of the biosensor is presented on the right. **(C)** Biosensors based on KB1691 or KB2123 detect Gαs activation by D1R. BRET responses were measured in HEK293T cells expressing D1R, Gβ1, Gγ2, Gαs-99V and either KB1691-Nluc or KB2123-Nluc. Results are the mean ± S.E.M. of n=3-5.

**Figure S2.**
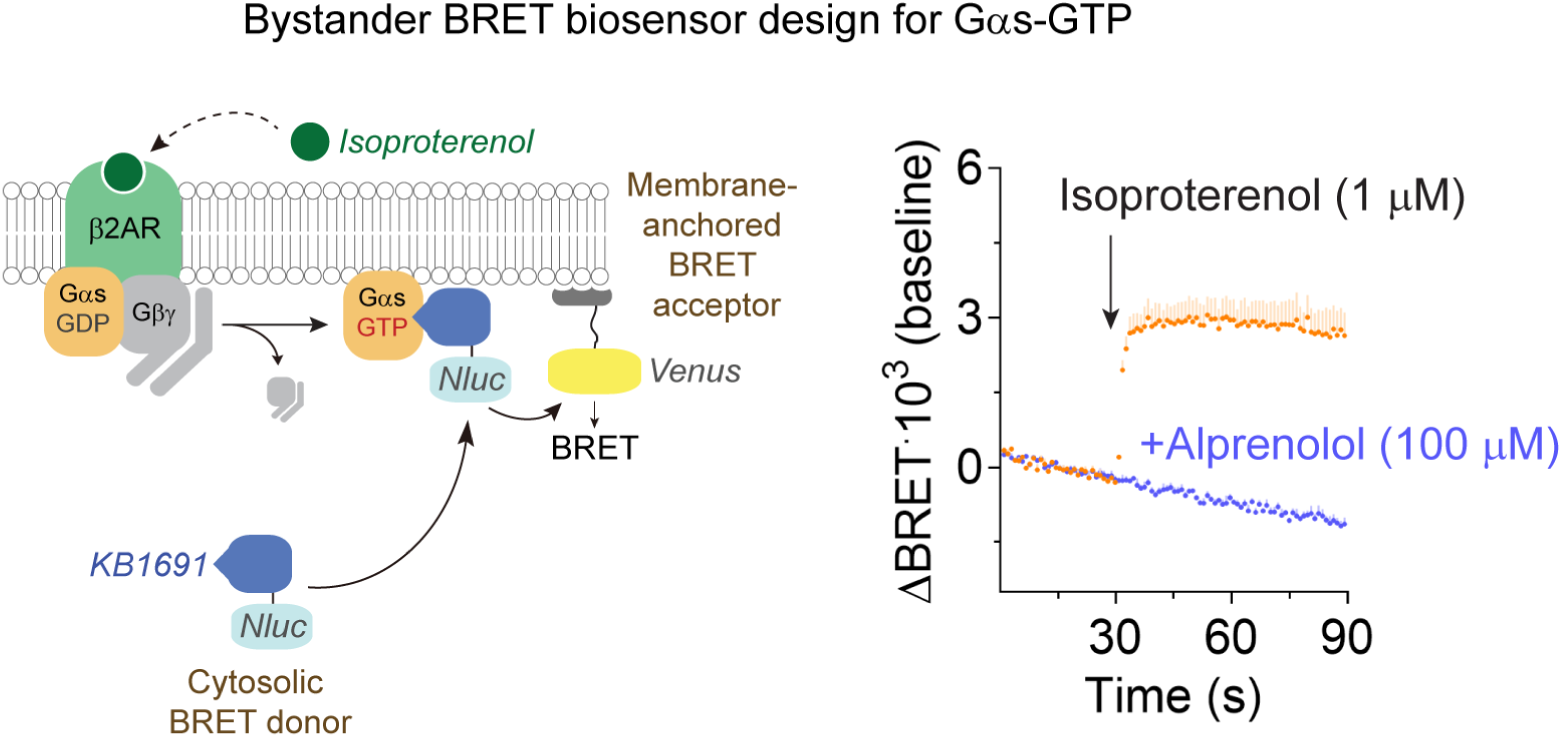
Bystander BRET biosensor design for active Gαs using KB1691-Nluc. *Left*, schematic representation of bystander BRET sensor system. KB1691-Nluc without any targeting sequence is expressed in the cytosol but is recruited to the plasma membrane upon GPCR-mediated activation of Gαs. A membrane-anchored Venus serves as the bystander BRET acceptor for KB1691-Nluc after translocation. *Right*, BRET was measured in HEK293T cells expressing the β2AR, untagged Gαs, cytosolic KB1691-Nluc, and a membrane anchored Venus. Results are the mean ± S.E.M. of n=4.

**Figure S3.**
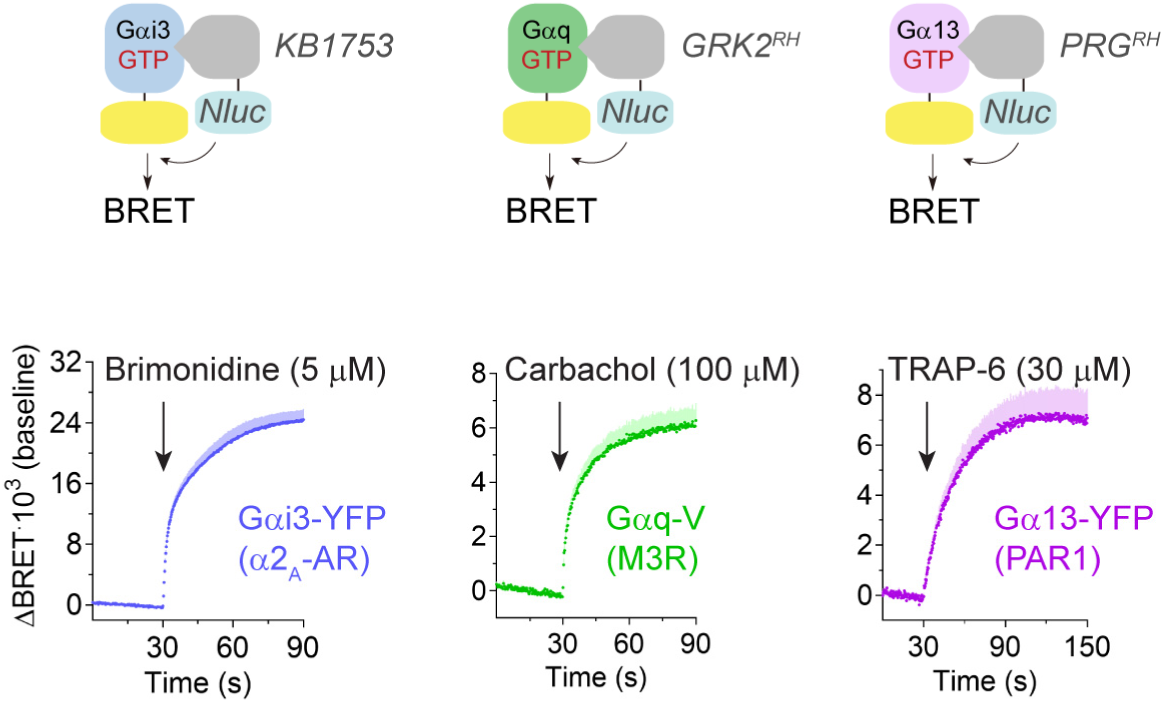
Detection of Gαi, Gαq and Gα13 activation with previously described biosensors. BRET responses were measured in HEK293T cells expressing the indicated combination of cognate GPCR, G protein, and Gα-GTP detector modules fused to Nluc. All Gα subunits were internally tagged with a YFP and they were co-transfected with Gβ1 and Gγ2. Experiments were done in parallel to those in Fig. 1C with the same stimulation conditions to serve as positive controls. Results are the mean ± S.E.M. of n=4-5.

**Figure S4.**
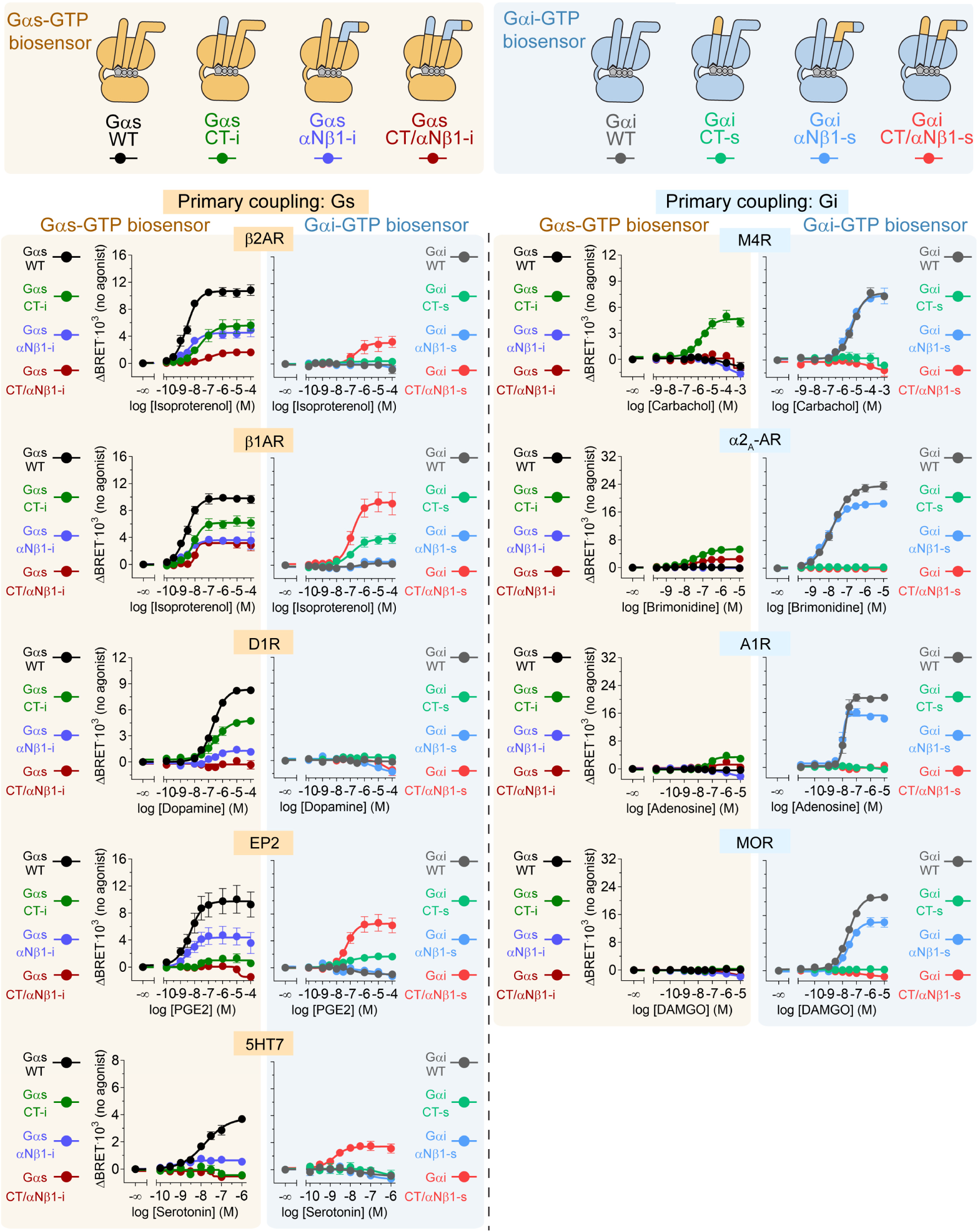
Contribution of the C-terminal tail (CT) and the αN/β1 ‘hinge’ of Gα subunits to their coupling to GPCRs. BRET was measured in HEK293T cells expressing the indicated combinations of GPCR, G protein and Gα-GTP detector modules fused to Nluc. All Gα subunits were internally tagged with a YFP and they were co-transfected with Gβ1, Gγ2, and either KB1691-Nluc (Gαs-GTP biosensor) or KB1753-Nluc (Gαi-GTP biosensor). Results are the mean ± S.E.M. of n=3-5.

**Figure S5.**
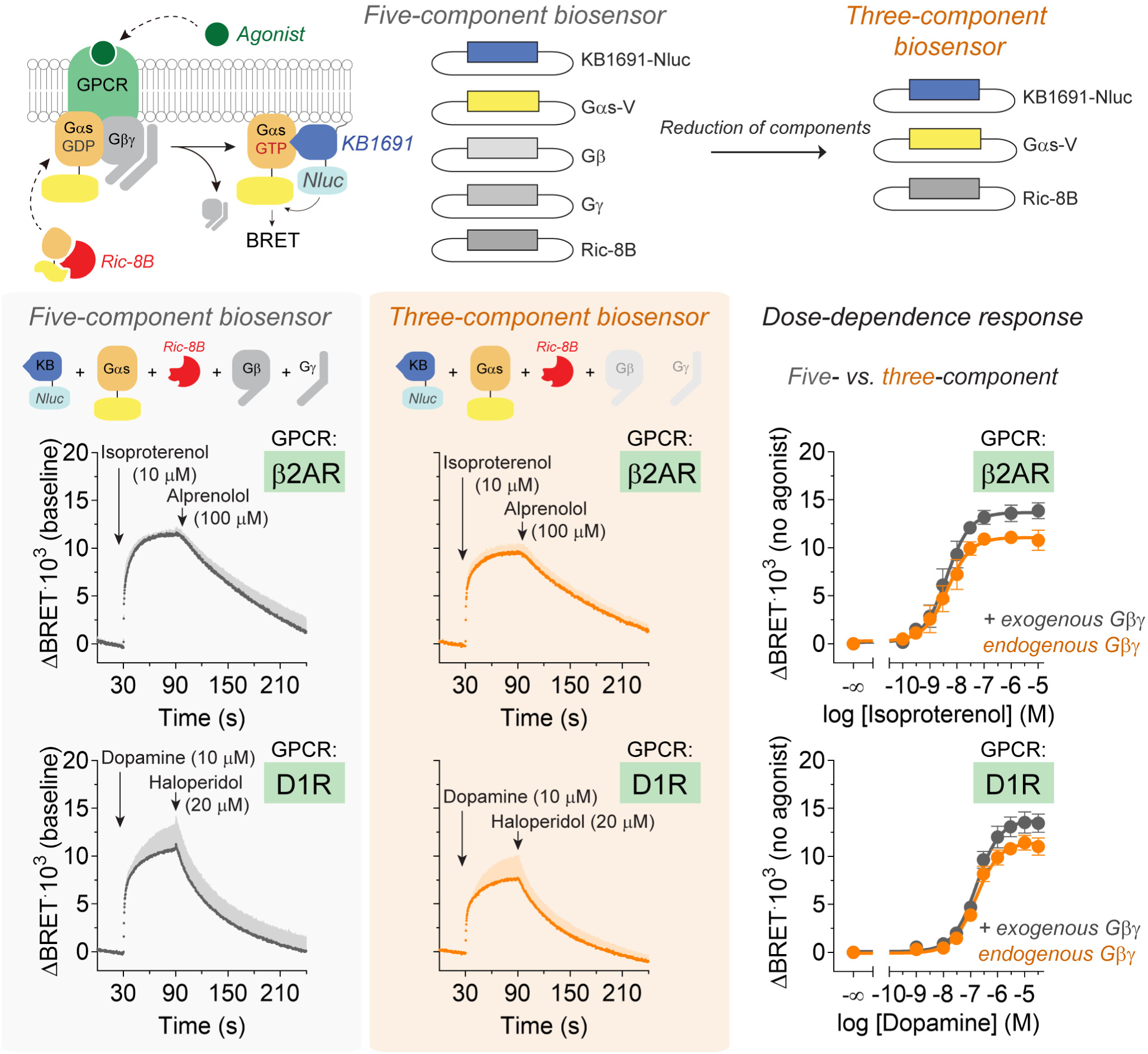
Comparison of Gαs biosensor performance with or without expression of exogenous Gβγ. *Top*, schematic of the reduction of Gαs-GTP biosensor components from five to three. *Bottom*, BRET was measured in HEK293T cells co-expressing KB1691-Nluc, Gαs-99V, Ric-8B, and the indicated GPCR (β2AR or D1R) with (black) or without (orange) exogenous Gβ1 and Gγ2. Results are the mean ± S.E.M. of n=3-5.

**Figure S6.**
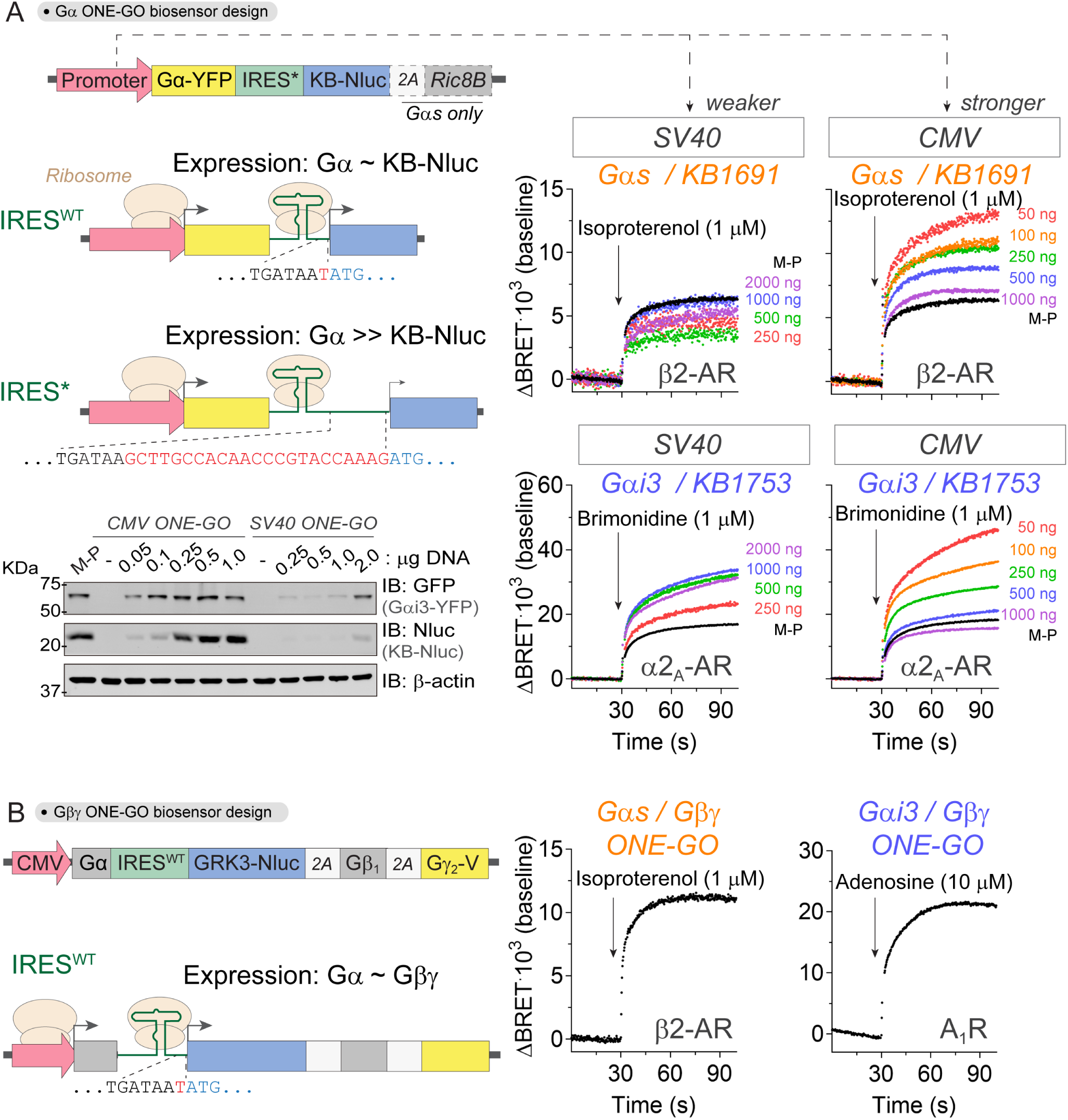
Optimization of single-plasmid ONE-GO biosensor designs for Gα-GTP or free Gβγ. **(A)** Gαs and Gαi3 ONE-GO biosensor designs. Two designs using a stronger or weaker promoter (CMV or SV40, respectively) were investigated for both Gαs and Gαi3 biosensors. In all cases, the Gα-YFP component was placed right after the promoter and the Nluc-fused component after a modified IRES (IRES*) of low efficiency to achieve higher acceptor-to-donor BRET ratios. BRET was measured in HEK293T cells transfected with different amounts of the indicated single-plasmid ONE-GO biosensors or their multi-plasmid (M-P) counterparts. β2AR or α2_A_-AR were co-expressed for Gαs or Gαi3, respectively. Results are the mean of n=2-3. A representative immunoblot showing expression of biosensor components is presented on the bottom left. **(B)** Gβγ ONE-GO biosensor designs. All constructs were based on a CMV promoter. Untagged Gα subunits were placed right after the promoter, followed by the Gβγ detector mas-GRK3ct-Nluc, Gβ1 and Venus tagged Gγ2 (separated by self-cleaving T2A sequences) after a high efficiency IRES (IRES^WT^). This favors the expression of Gα and Gβγ at similar level to facilitate their assembly into a functional heterotrimer. BRET was measured in HEK293T cells expressing the indicated combination of Gα/Gβγ ONE-GO biosensor and GPCR. Results are representative traces of an individual experiment.

**Figure S7.**
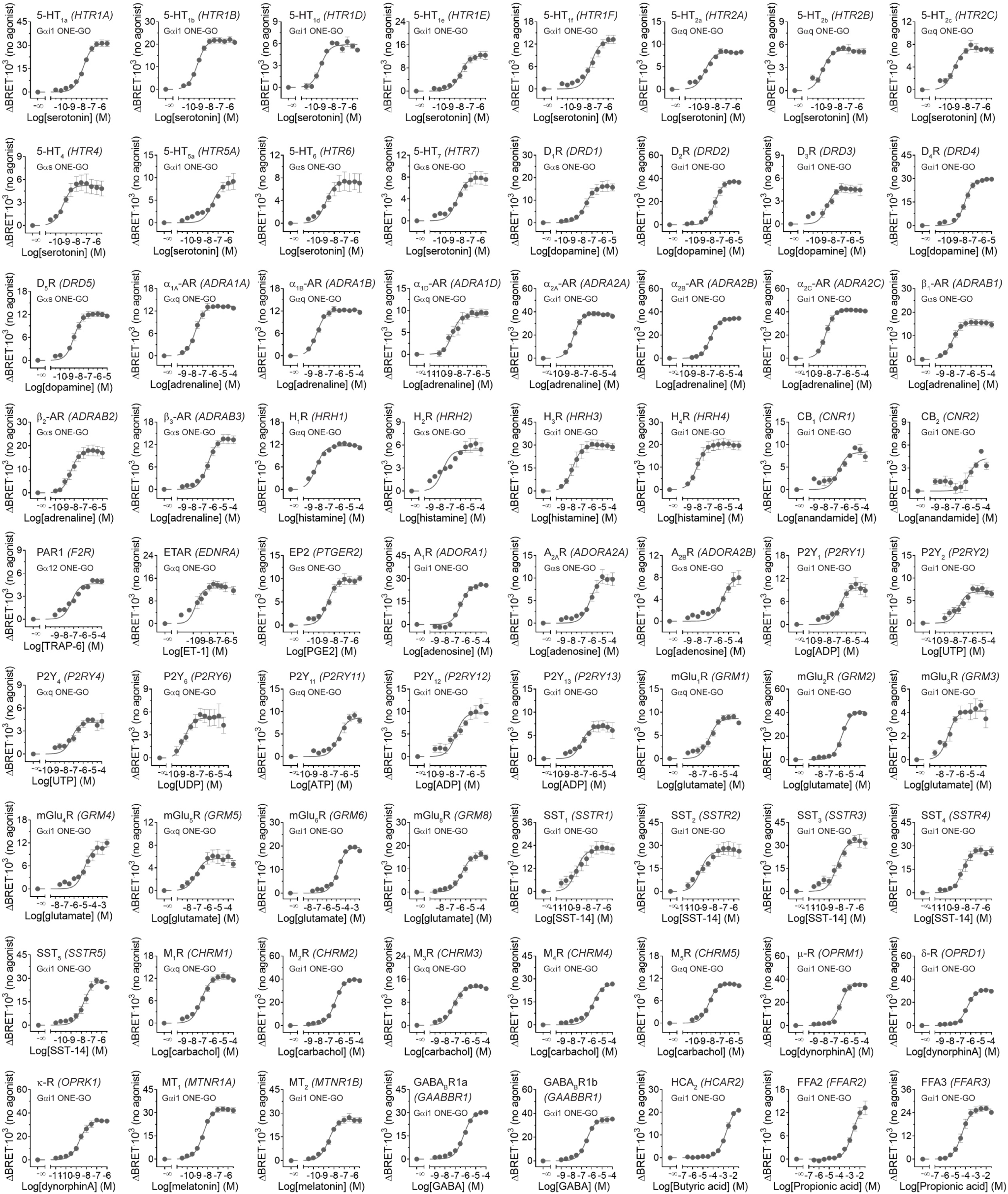
ONE-GO biosensors report the activation of many GPCRs. BRET was measured in HEK293T cells expressing the indicated combinations of GPCR and ONE-GO biosensor upon stimulation with the indicated agonists. Results are the mean ± S.E.M. of n=3-5.

**Figure S8.**
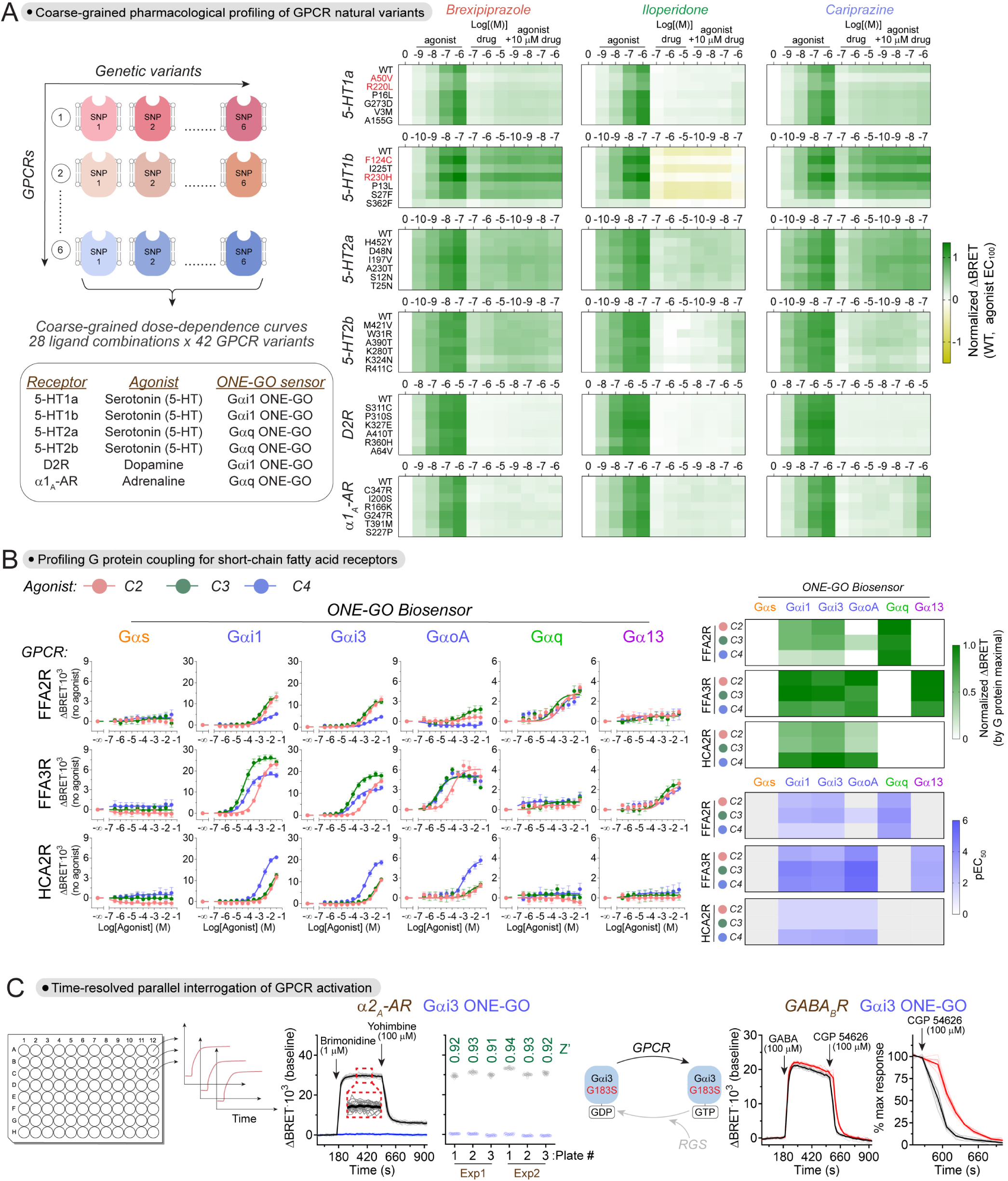
Large-scale parallel interrogation of GPCR activity with ONE-GO biosensors. **(A)** Pharmacogenomic profiles of atypical antipsychotics. *Left*, schematic of the assay. *Right*, BRET responses were measured in HEK293T cells expressing the indicated GPCRs and ONE-GO biosensors upon stimulation with the indicated concentrations of antipsychotic drug, of cognate agonist, or a combination of both, as shown. Results are the mean of n=3-7. **(B)** Profiling of G protein selectivity across short-chain fatty acid receptors. *Left,* BRET responses were measured in HEK293T cells expressing the indicated combination of GPCR and ONE-GO biosensor upon stimulation with different short chain fatty acids, as shown. Results are the mean ± S.E.M. of n=4-6. *Right*, heatmaps represent maximal responses normalized by biosensor and pEC_50_ values calculated from dose dependence curve fits presented on the left. **(C)** Time-resolved parallel interrogation of GPCR responses. *Left*, schematic of the assay. *Middle*, BRET was measured in HEK293T cells expressing the α2_A_-AR and the Gαi3 ONE-GO biosensor, which were treated with ligands as indicated. The graph on the left is a representative experiment of 48 simultaneously kinetic measurements (thin traces are individual wells and the thick line is the average). The graph on the right is a scatter plot of Z’ values of 3 individual 48-condition recordings (i.e., in 3 plates) repeated in 2 independent experiments. *Right*, BRET was measured in HEK293T cells expressing the GABA_B_R and ONE-GO biosensor for either Gαi3 (black traces) or for Gαi3 G183S (red traces). Results are 24 kinetic measurements for each G protein carried out simultaneously (thin traces are individual wells and the thick line is the average) from one representative experiment of 3.

**Figure S9.**
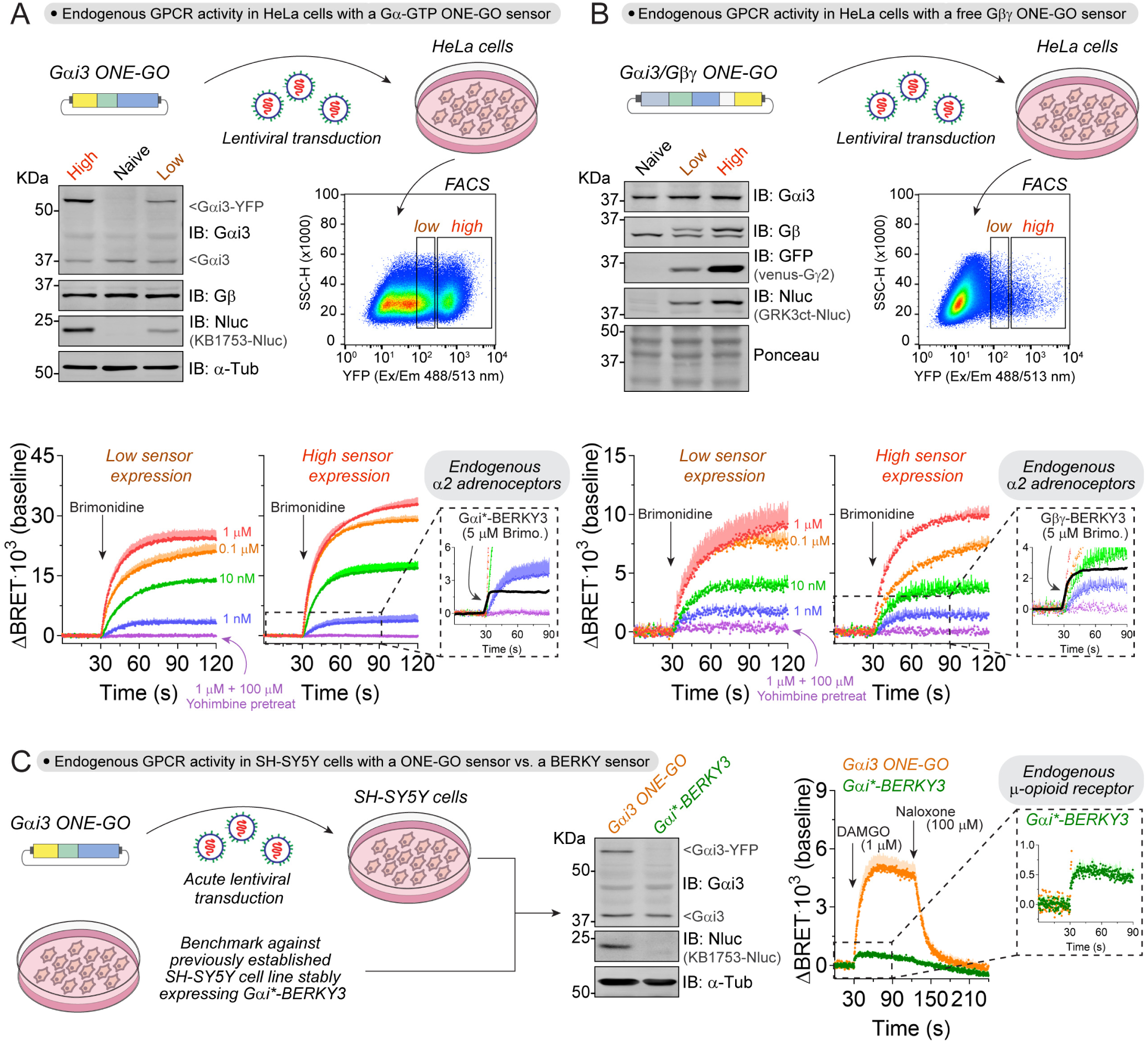
Detection of endogenous GPCR activity in cell lines with ONE-GO biosensors. **(A-B)** Detection of endogenous GPCR activity in HeLa cells stably expressing ONE-GO biosensors. *Top*, generation of HeLa cells stably expressing high or low levels of either Gαi3 ONE-GO **(A)** or Gαi3/Gβγ ONE-GO **(B)** biosensors by lentiviral transduction followed by FACS. A representative immunoblotting result to confirm the relative expression of biosensor components and endogenous G proteins is shown on the bottom left. *Bottom*, BRET was measured in HeLa cells stably expressing either the Gαi3 ONE-GO or the Gαi3/Gβγ ONE-GO sensor, and treated with ligands as indicated. The black traces in the insets correspond to previously described^30^ BRET responses detected in HeLa cells expressing another biosensor for the same G proteins (Gαi*-BERKY3 in **A** and Gβγ-BERKY3 in **B**) upon stimulation of endogenous α2 adrenergic receptors. Results are the mean ± S.E.M. of n=2-3. **(C)** Detection of endogenous opioid receptor activation in SH-SY5Y cells acutely transduced with a ONE-GO biosensor. BRET was measured in parallel either in SH-SY5Y cells acutely transduced with a lentivirus for the expression of Gαi3 ONE-GO biosensor (orange), or in a previously established^30^ SH-SY5Y line stably expressing the Gαi*-BERKY3 biosensor (green) upon stimulation with the μ-opioid receptor agonist DAMGO. Results are the mean ± S.E.M. of n=4. The immunoblot shows the expression of ONE-GO biosensor components and endogenous Gαi3.

**Figure S10.**
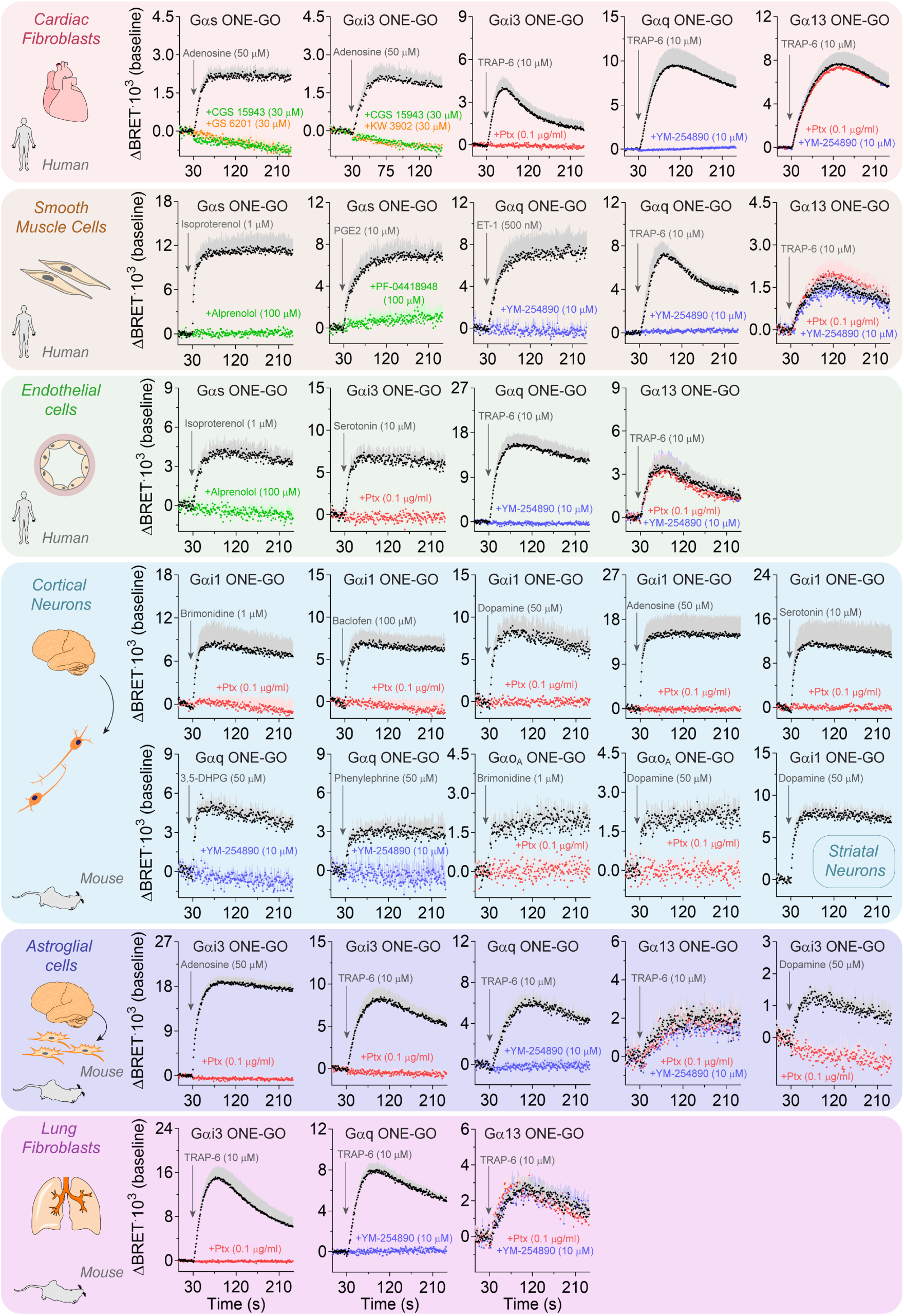
Endogenous GPCR responses and respective controls detected with ONE-GO biosensors in various primary cell types. BRET was measured in the indicated cell types after acute transduction with lentiviruses for the expression of different ONE-GO biosensors, as indicated. Cells were stimulated with agonists as indicated during the BRET recordings. In some cases, cells were pre-treated with the indicated GPCR antagonists (green and orange) or the G_q/11_ inhibitor YM-254890 (blue) for two minutes before initiating the measurements, or with pertussis toxin (Ptx, red) overnight. Results are the mean ± S.E.M. of n=4-6.

